# Forests and woodlands resistant to drought revealed in remotely sensed foliar moisture content using probabilistic models

**DOI:** 10.64898/2025.12.10.692240

**Authors:** Ivan Kotzur, Ben D. Moore, Matthias Boer, Marta Yebra, Kara Youngentob

## Abstract

Foliar moisture content (FMC) is of primary importance in many biological and ecological processes of forests and woodlands, including herbivory and fire dynamics. The role of FMC, which is the mass of moisture on a dry matter basis, in such processes is increasingly important due to changes in length and severity of climate extremes, like drought. Here we studied the sensitivity of forest and woodland FMC to climatic water balance, i.e. Standardised Precipitation and Evapotranspiration Index (SPEI), across five landscapes in temperate and sub-tropical south-eastern Australia. To do this we used copula-probabilistic modelling, including consideration of drought (i.e. SPEI) length. We used an FMC dataset over eight years of Sentinel-2 satellite reflectance (2015-2023), based on the inversion of a radiative transfer model at 20m resolution. We found the spatial variation of FMC within landscapes to be large and that the spatial variation in response to drought was very large, particularly within the more arid landscapes. The areas of lesser response to extreme drought are revealed to be drought resistant forests and woodlands that are potential climate refugia for animals, and natural breaks in the contiguity of fire fuels.

## 1. Introduction

The response of foliar moisture content (FMC) in forests and woodlands to drought is very important because FMC is crucial to plant metabolism and growth, which in turn affects ecosystem water balance, productivity and carbon cycles. Indeed, leaf water content may be the primary plant trait (Wang et al. 2022), and this importance is reflected in how climatic water gradients control the transitions between ecosystems across landscapes (e.g. woodlands to savanna, Williams, Shuman and Bartlein, 2009). Considering the importance of FMC, its response to drought will influence the potential for forests to support herbivory, via changes to forage quality and quantity (Briscoe et al. 2016, Gely et al. 2020), and the response will influence forest fire risk (Rao et al. 2022) because fuel availability is strongly affected by plant biomass and dryness (Nolan et al. 2016). These ecological dynamics are particularly important in high temperatures when critical thresholds of animal hydration and fire behaviour may be approached (Cahill et al. 2013, Choat et al. 2018). Such thresholds may be due to sudden FMC declines, there have been rapid canopy browning events during recent droughts (Losso et al. 2022). In some ecosystems such conditions have led to large mortality events in arboreal mammal populations (e.g. Lunney et al. 2012, Gordon et al. 1988). These biological responses are being driven by climate extremes, including higher temperatures, greater temperature extremes, and more severe droughts, with the latter two phenomenon becoming more regular in future (Zhongming et al. 2021). These ‘global change-type droughts’ are hotter with higher evapotranspiration from ecosystems increasing water loss (Allen et al. 2015). During droughts the supply of water to trees from the soil is reduced as soil moisture declines, while in high temperatures the vapour pressure deficit of the atmosphere increases, demanding more moisture from trees via leaves (Griebel et al. 2023, Lyons et al. 2021). Such environmental variations affect animals directly via physiological stress from increased air temperature, and indirectly, via declining habitat quality with reduced leaf moisture and productivity (Cahill et al. 2013). These environmental effects may change at different rates across space and time due to landscape, soil and microenvironmental factors, potentially creating climate refugia for forest animals (Keppel et al. 2012). Therefore, the response of vegetation in species’ habitats to climate extremes like drought is ecologically important (Rosenblatt & Schmitz 2016). Furthermore, the present and future variation in FMC of forests and woodlands may be large (Konings et al. 2021), understanding these patterns is important in animal and fire ecology (Griebel et al. 2023, Briscoe et al. 2016).

FMC is the weight of water in vegetation as a percentage of the dry weight of that vegetation.

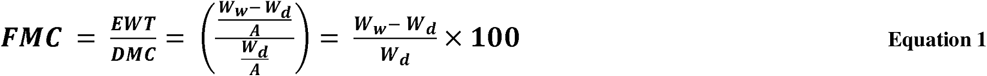

Where the equivalent water thickness (EWT) is the water mass per unit leaf area, dry matter content (DMC) is the dry weight per unit leaf area, W_W_ is the fresh or wet weight of vegetation, W_d_ is dry weight and A is leaf area. The conversion of FMC on a dry weight basis to water content (WC) on a wet weight basis (i.e. fresh weight) produces a metric that may be more intuitive in animal ecology because it informs how much pre-formed water the animal ingests per wet mass of foliage:

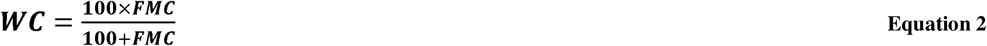

FMC is an important predictor of fire activity (Griebel et al. 2023) and the variable has been analysed mostly in that context including in remote sensing methodologies (Gale et al. 2021, Kotzur et al. 2025). However, it is a measure of vegetation water and is equivalent to many similar measures of plant composition made on a dry weight basis (e.g. foliage quality, Beale et al. 2023), making it applicable outside fire ecology.

Climatic drought occurs when rainfall is less than normal in an area, whereas ecological drought occurs when plants are affected by such water deficit. Ecological drought influences FMC variation in trees, with increased environmental moisture deficit generally driving FMC downward (Nolan et al. 2022). The magnitude of this response is determined by tree physiology including hydraulic vessel structures and functions, supply of water to trees from the soil and demand for water by the atmosphere (Brodrick et al. 2019). Trees adjust water balance to maintain hydraulic function (Bartlett et al. 2012), though in severe drought the capacity of this adjustment can be exceeded resulting in physiological damage to hydraulic vessels (Choat et al. 2018). Water supply to trees is reduced by declining precipitation and subsequently soil moisture, these processes being mediated by topography and soil characteristics (Anderegg et al. 2013). These landscape influences are relatively static at the timescale of droughts. Additionally, spatially explicit measures of vegetation response incorporate topographic and soil variation, particularly if using gridded data in which the spatial resolution is approaching scales of landscape variation (e.g. moderately fine scale remote sensing data). Water demand by the atmosphere from vegetation is determined by vapour pressure deficit (VPD), sunlight and temperature (Paz-Kagan & Asner 2017). It is therefore important to consider temperature (i.e. basis of VPD) in any measure of ecological drought, rather than considering water supply alone (Vicente-Serrano et al. 2010).

As well as affecting FMC, drought is a condition in which FMC may be of increased importance for animal habitat quality. When climatic water is in deficit and tree water balance is affected, it is expected that FMC decline could alter habitat quality (Briscoe et al. 2016). This concerns specialist, mammalian folivores particularly, for which low FMC may be an insufficient water supply for thermoregulation, unlike in most conditions (Moore et al. 2004). Thereby droughts, particularly extreme droughts, may reduce vegetation water availability to critical levels (Melzer 1994, Hume & Esson 1993), during a time when open water sources are likely to have diminished (e.g. tree surfaces, creeks, Mella et al. 2019). Furthermore, droughts often involve increased temperature and coincide with warm seasons in many biomes (Spinoni et al. 2014). High temperatures increase the use of water in animal thermoregulation and so water demand by the animal (Turner 2020, Beale et al. 2018). Additionally, temperature extremes (i.e. heatwaves) can cause FMC to rapidly decline (Fox-Hughes et al. 2021). Drought effects on FMC may also impact whole food webs via insects, particularly through those leaf-feeding and sap-sucking species which rely on nitrogen-rich compounds made available by leaf turgor pressure which declines with moisture content (Gely et al. 2020). The co-occurrence of elevated water demand in animals and dehydrated tree canopies will impact habitat quality and potentially survival (Briscoe et al. 2016).

Brodrick et al. (2019) modelled remotely sensed canopy water content of forests as an indicator of drought resistance across landscapes in the state of California (USA), by quantifying the integral of a drought index when canopy water is declining through a moderately below-average range before passing statistical thresholds (i.e. drought affected). The study showed that most forests are less resistant to long, severe droughts, though there was substantial variation in drought resistance amongst forests. This approach successfully highlighted areas where vegetation water is maintained. However, no such study has been completed for ecosystems in Australia, in which response strategies to drought may differ from those in USA, including the ability of many common *Eucalyptus* species to resprout from epicormic buds (Zeppel et al. 2015).

Analysing FMC together with drought presents intrinsic and methodological challenges. As shown earlier FMC is dependent on climatic water and therefore covaries with drought. This dependence is likely to be highly variable across space owing to differing environmental and species responses to drought (Paz-Kagan & Asner 2017). Copula modelling can account for this by considering marginal distributions of variables and a joint distribution (i.e. dependence structure) separately (Favre et al. 2004). The joint distribution may be represented by one of several copula functions that represent different dependence structures (Anderson et al. 2019), which may be applied to gridded data across space. The variables can be represented by any type of differing marginal distributions including non-normal distributions. FMC values in forests during drought likely make up the tail of a positively skewed FMC distribution (i.e. non-normal). Furthermore, study of rare phenomenon, for example low FMC and extreme drought, inherently involves few data points. This can partly be overcome by a regular data collection over a long timeseries (e.g. satellite remote sensing), and by a copula modelling framework which evaluates extreme exceedance in the joint probability distribution function rather than frequency of critical extreme values in the random variables themselves (Zscheischler & Seneviratne 2017). For animal species adapted to a range of environments, the critical value of habitat resources may be difficult to generalise (e.g. FMC, Hume and Esson, 1993; Melzer, 1994). This challenge is exacerbated by the different degrees of tree adaptations to drought across habitats (Nolan et al. 2016, Briscoe et al. 2015). These issues are partly allowed for by analysing vegetation response as a timeseries percentile (or set of percentiles) because it indicates the value relative to the distribution of each pixel, accounting for different mean values across space. Percentiles are integral to a copula framework (Guo et al. 2023) and have also been used in studies of vegetation sensitivity to drought (Konings et al. 2017).

In this study we monitored FMC using remote sensing over the period 2015-2023 in five landscapes in temperate and subtropical forests and woodlands in the state of New South Wales (Australia). This included a very wet and a very dry period, in which extremely low levels of vegetation water were involved in large bushfires (Boer et al. 2020, Fox-Hughes et al. 2021). Such events can impact animal occupancy particularly in combination with drought (Lunney et al. 2017). The aim of the study was to identify forests and woodlands where high FMC tends to persist through time, using a probabilistic approach. To do this we assessed forest FMC variation and its relationship with climatic drought in terms of strength, temporality, and likelihood of decline in extreme drought, both across and within landscapes. To this end we posed the following questions: 1). How does FMC vary across and within forests and woodlands? We hypothesised that high FMC landscapes and tree pixels will be more stable through time (H_1_) because of less variability in water supply. 2). To what degree does climatic drought impact FMC and over what timescales? We expected there to be a greater degree and more rapid impact in regions of moderate aridity compared to areas of high aridity (H_2_) due to differences in trees tolerance of drought and intensity of drought experienced. 3.) How likely are forests and woodlands to decline to low levels of FMC in drought? And where is the variation in these probabilities throughout landscapes? Our hypotheses were that the general likelihood would be low because drought is relatively rare (H_3_), however that there will be great contrasts in likelihood within landscapes at the scale of hundreds of meters (H_4_) due to topography, soils and other micro-environmental gradients.

## 2. Methods

### 2.1 Study areas

To highlight within– and between-landscape variability in FMC response to drought we focused on 5 landscapes with an area of 10 km^2^ each. We selected these landscapes to capture a range climatic conditions and vegetation structural types, in the state of New South Wales, Australia (Figure 1, Figure 2, Table 1). We selected landscapes with little history of fire in the two years before or anytime during the study period, to minimise the confounding effects of fire and subsequent vegetation recovery on canopy condition, as well as the need to discard data. Given the prevalence of fire in the study region, we were not able to find five landscapes of interest with no fire at all. The landscapes mostly comprised National Parks and Nature Reserves, and in one case, Military Reserve (i.e. Kentlyn). Climate ranged from temperate to sub-tropical, with precipitation generally declining with distance from the coast.

**Figure 1.**
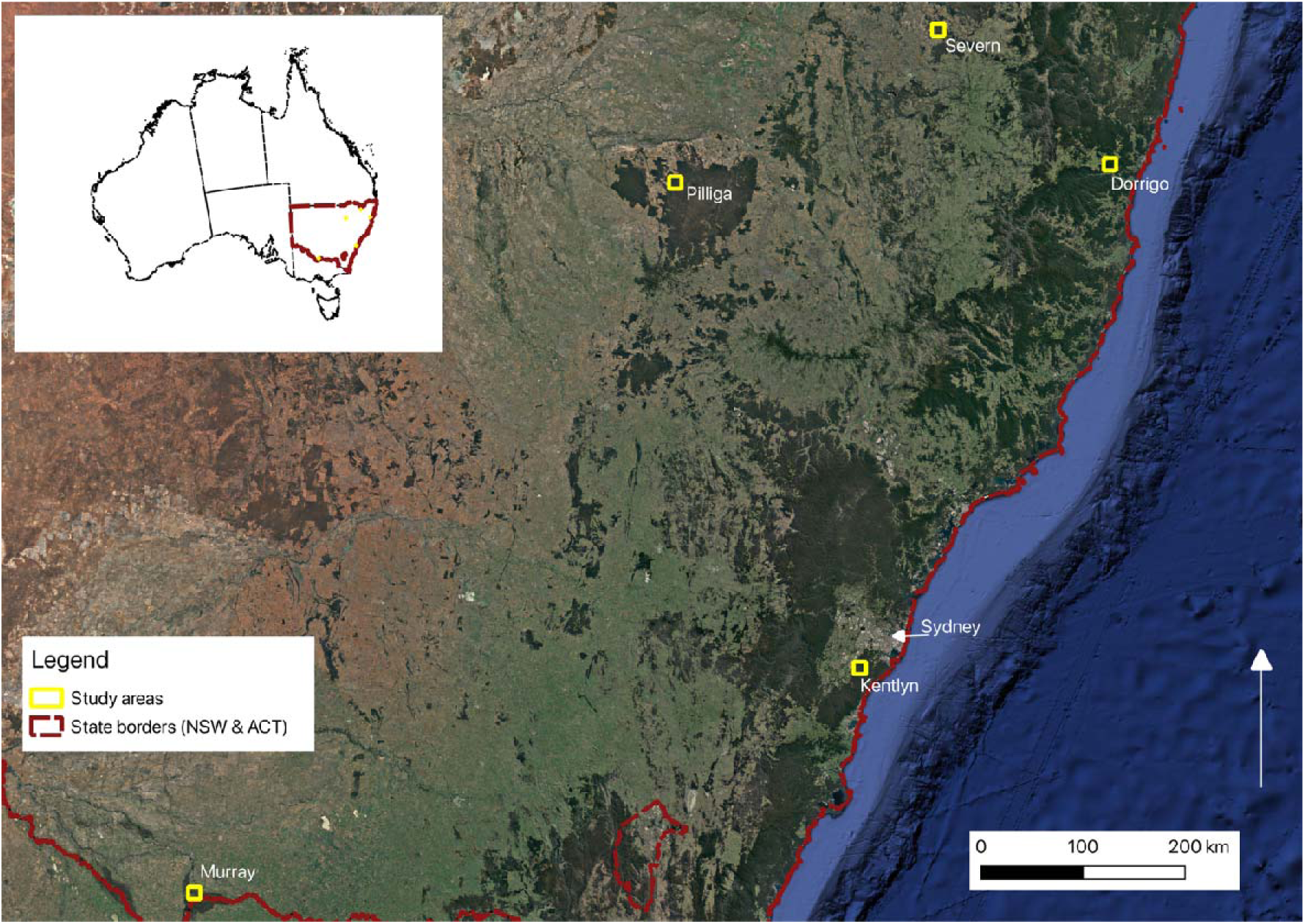
Locations of study areas in temperate and sub-tropical forests and woodlands in the state of New South Wales (Australia [inset map]). Study landscapes names from west to east: Murray, Pilliga, Kentlyn, Severn (northern most), Dorrigo.

**Figure 2.**
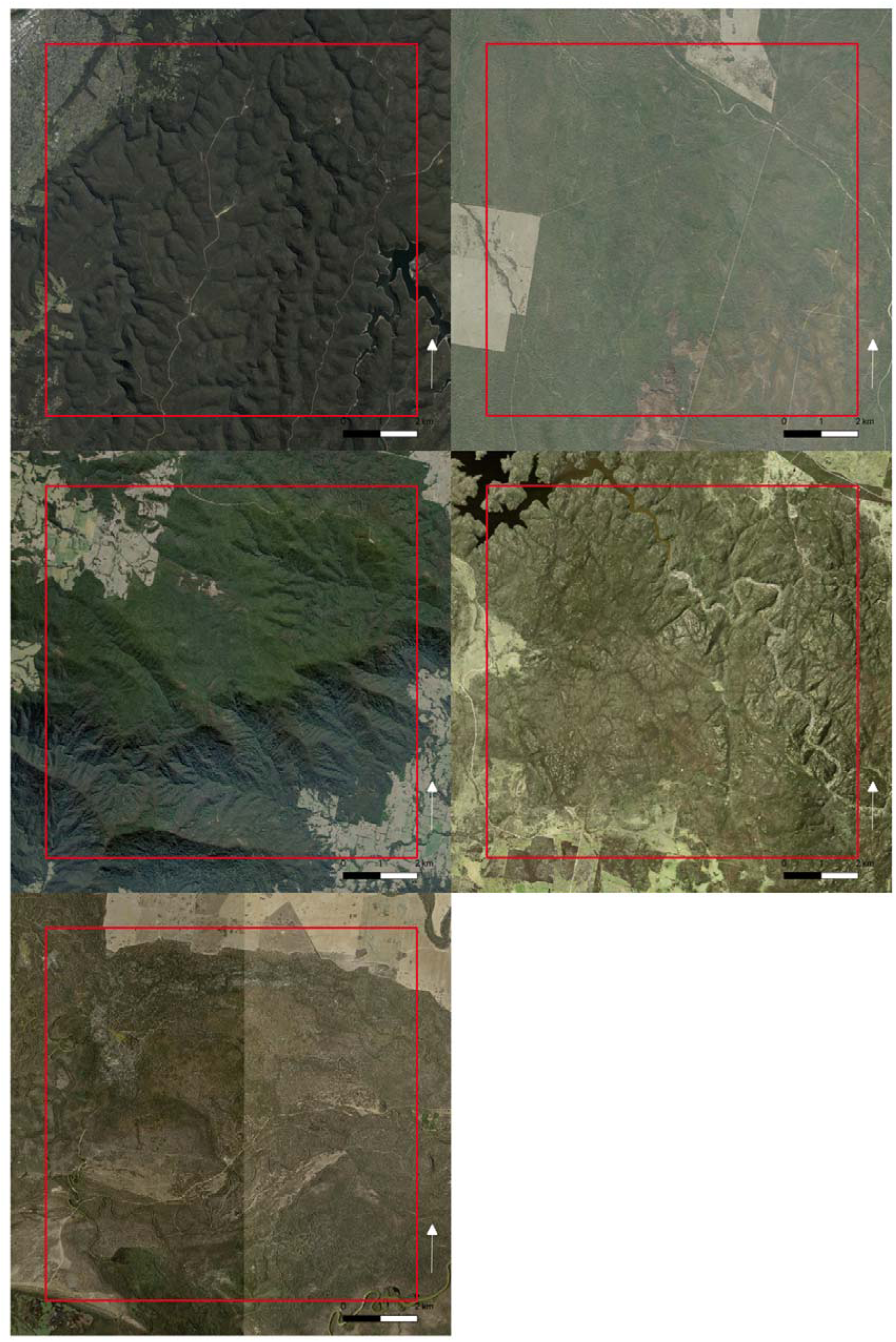
Satellite imagery of study landscapes, each being 10×10km in area. Landscape names from left to right, top to bottom: Kentlyn, Pilliga, Dorrigo, Severn, Murray. Larger version of images available, see link in Data. Availability below. Imagery from various dates, source: https://maps.six.nsw.gov.au/arcgis/rest/services/public/NSW_Imagery/MapServer.

**Table 1.** Locations and vegetation types of study areas in the state of New South Wales (Australia). Plant community.

| **Study area** | **Plant community type - predominant** | **Location (centre)** | **MAP (mm)** | **MAT (°C)** | **Elevation range (m)** |
| --- | --- | --- | --- | --- | --- |
| **Kentlyn** | *(Dry Sclerophyll Forests (Shrubby sub-formation))*<br>*Scribbly Gum Woodland / Apple-Blackbutt Gully Forest / Sandstone Gully Forest* | -34.10599°,<br>150.88038° | 966 | 21.4 | 42 – 292 |
| **Pilliga** | *(Dry Sclerophyll Forests (Shrubby sub-formation))*<br>*Narrow-leaved Ironbark - White Cypress Pine - Buloke tall open forest* | -30.60629°,<br>149.17498° | 620 | 25.9 | 213 – 274 |
| **Dorrigo** | *(Rainforests) Coachwood Warm Temperate Rainforest* | -30.35103°,<br>152.82256° | 1906 | 19.1 | 18 – 1037 |
| **Severn** | *(Dry Sclerophyll Forests (Shrubby sub-formation))*<br>*Tumbledown Gum-Ironbark Exposed Forest* | -29.43463°,<br>151.33504° | 777 | 22.5 | 503 – 776 |
| **Murray** | *(Forested Wetlands) River Red Gum herbaceous-grassy very tall open forest wetland on inner floodplains* | -35.78993°,<br>145.00879° | 390 | 22.6 | 83 – 116 |

type maps produced in OEH (2017).

### 2.2 Foliar moisture content modelling

To remotely sense FMC across the study areas we used inverse radiative transfer modelling (RTM) based on reflectance data from the Multispectral Instrument onboard the two Sentinel-2 satellites (Yebra et al. 2008, Jurdao et al. 2013). The method is described in detail elsewhere (Kotzur et al. 2025). Briefly, an RTM includes forward modelling to simulate the reflectance of leaves and canopies based on biophysical principles and vegetation characteristics, and inverse modelling which uses simulated reflectance to link the characteristics of interest (FMC in this case) to observed reflectance in the study area. From the published forward modelling we filtered the resultant look-up-table of simulations, to vegetation type and the leaf area domain of interest and transformed it to match wavelengths of the Sentinel-2 sensor (Table 2). We inverted the RTM (LUT) using the spectral angle function and optimised this process based on a validation with field FMC samples across forests and woodlands of NSW. This resulted in an acceptably accurate model (r^2^=0.62, *P<*0.01, RMSE=19.92% dry matter content) relative to previous inverse radiative transfer models of the same variable in forests (r^2^=0.17, *P<*0.01, RMSE=32%, (Yebra et al. 2018). To efficiently scale up the application of the foliar moisture model from site-pixels to landscapes we then used an emulation approach (see Kotzur et al. 2025).

**Table 2.** Spatial data inputs for regression and probabilistic modelling of foliar moisture with drought over 5 landscapes in forests of New South Wales, including data for masking of non-forest.

| Source | Variable | Study use | Native spatial resolution | Native temporal resolution |
| --- | --- | --- | --- | --- |
| **Remotely sensed radiative transfer model**<br>(Kotzur et al. 2025) | Foliar moisture content (FMC, % weight dry matter) | Response variable | 20 m | 5 day<br>(dependant on satellite reflectance data quality) |
| **SILO Climate data**<br>(Jeffrey et al. 2001, Zajackowski & Jeffrey 2020) | Rainfall (mm)<br>Potential evapotranspiration (mm) | Input for drought index | 0.05° (~5km) | Daily |
| **Foliage Projected Cover NSW**<br>(Fisher et al. 2016) | Canopy density above 2 m | Vegetation mask | 5m | Instantaneous<br>(various acquisition dates 2008-2011) |
| **Native Vegetation Extent NSW**<br>(Fisher et al. 2016) | Native vegetation |  |  |  |
| **NSW State Vegetation Type Map**<br>(Roff et al. 2022) | Plant community types | Vegetation mask | 5m | Instantaneous<br>(various acquisition dates) |
| **GLAD Forest Canopy Height**<br>(Potapov et al. 2021) | Forest height | Vegetation mask | 25m | Instantaneous<br>(2019 acquisition) |
| **Soil and Landscape Grid of Australia** (Searle et al. 2022) | Topography | Predictor variables | 30m | Instantaneous<br>(various acquisition dates) |

### 2.3 Data sources

### 2.4 Fire scar masking

To minimise the effect of fire on the analysis we masked out pixels in the FMC data based on a fire history dataset, which is updated periodically (DPIE 2010). This spatial data is regularly updated and contains a polygon for each fire and the date it started. We selected fires which intersected with each landscape only if they occurred during or in the two years preceding our FMC observations, then converted these polygons to rasters matching the FMC spatial resolution and coordinate system. We chose a period of 2 years as the duration after fire in which the FMC may be affected. (Khanal et al. 2014) found that within this duration, the satellite reflectance of more than 90% of fire-affected forest pixels had recovered, in terms of satellite reflectance. Based on this length of time, we extended the fire mask rasters in the time dimension. The 3D mask was then applied to the input data, retaining any pixels outside the space or time of the fire affected period.

### 2.5 Drought analysis

To quantify the timing of the response of FMC to drought we tested correlations with the Standardised Precipitation Evapotranspiration Index (SPEI, Vicente-Serrano et al. 2010) at different time scales (i.e. persistence of water deficit). The Standardised Precipitation Evapotranspiration Index is based upon precipitation and temperature, via potential evapotranspiration (PET) and indicates deviations from the water balance and includes the fitting of log-logistic probability distributions over the chosen time scale or scales (Vicente-Serrano et al. 2010). To allow for a wide range of forest FMC responses we chose SPEI timescales of 1 to 48 months, using climate data from *SILO* (not an acronym, Table 3). In this way different drought effect response times may be analysed by the same metric and accounting for cumulative and climate change influences (Vicente-Serrano et al. 2010). We completed a correlation analysis of the FMC and SPEI time series per pixel, for every timescale of SPEI (i.e. 1 to 48 months), with the highest Pearson’s R correlation coefficient taken to indicate the water balance variation with the greatest effect on FMC for each vegetation pixel. This resulted in three maps, one indicating the highest degree of water balance variation effect on FMC in general rather than that for one given length of drought (i.e. negative variation). The second resulting layer was of the significance of the former FMC-SPEI relationships. Thirdly we extracted the timescale index of the strongest FMC-SPEI relationships (i.e. first result), which was a layer of the number of months at which water balance variations are seen most strongly in FMC.

**Table 3.** Summary statistics of foliar moisture content of forests and woodlands at 5 landscapes in temperate and sub-.

| **Landscape** | **Foliar moisture content (% dry matter content)** |  |  |  |  |  |  |  |  |
| --- | --- | --- | --- | --- | --- | --- | --- | --- | --- |
|  | **Mean** | **Std.** | **Min.** | **Max.** | **Median** | **Variance** | **Skewness** | **Kurtosis** | **n** |
| **Severn** | 79.01 | 30.37 | 36.93 | 204.71 | 66.21 | 922.28 | 0.84 | -0.51 | 62514967 |
| **Pilliga** | 80.88 | 31.71 | 37.15 | 192.93 | 68.98 | 1005.59 | 0.79 | -0.59 | 67297209 |
| **Murray** | 120.53 | 28.21 | 37.65 | 207.03 | 127.95 | 795.60 | -1.09 | 0.25 | 28596444 |
| **Kentlyn** | 125.74 | 20.04 | 37.18 | 208.80 | 129.80 | 401.61 | -1.18 | 1.85 | 54183795 |
| **Dorrigo** | 141.97 | 9.11 | 38.65 | 213.11 | 143.77 | 83.02 | 0.26 | 13.57 | 54589041 |

### 2.6 Conditional probabilities analysis

We used copula functions to model the dependence structure (Nelsen 2006) of FMC and drought, per pixel. Copula functions are multivariate cumulative distribution functions which describe the interrelations of random variables. We used the bivariate form:

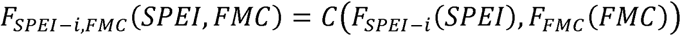

where *SPEl* – *i*, is the SPEI timescale of the interval (*i*) that is most strongly correlated with FMC at each pixel. *C*() is a copula function. *F_SPEI-i,FMC_* is the joint distribution, *F_SPEI-i_* and *F_FMC_* are marginal distributions of the *SPEIi* and FMC timeseries, respectively. By design, the drought index, SPEI, is normally distributed. For the response variable, FMC, we identified the marginal distribution by fitting many possible distributions, available in the python package SciPy (Virtanen et al. 2020), to the data, by calculating the sum of squares error and the Kolmogorov-Smirnov test, as well as visualising the probability distribution function against histograms of subsamples of the data (i.e. random pixels). These subsamples were taken from across space and time, and the two dimensions combined. The final set of distributions used in the copula models included generalised normal, generalised logistic, logarithm gamma, normal inverse Gaussian, Johnson SU, Johnson SB, Burr (Type III), Cauchy and folded Cauchy distributions.

We used the python package Pyvinecopulib to fit the copula models to the CDF-transformed marginal variables based on maximum likelihood estimation (Nagler & Vatter 2019). Copula families tested included Gausssian, Student, Clayton, Gumbel, Frank and Joe (Nagler & Vatter 2019, Nelsen 2006, Joe 1997). We compared the goodness of fit for each family, using the Akaike information criterion (AIC), and we considered the optimal copula to be that with the lowest AIC score.

To estimate the likelihood of FMC decline during drought we evaluated the probability distribution function of the fitted copula for a set of conditions, following (Fang et al. 2019). The conditional variable was SPEI and for each tested drought condition (i.e. SPEI level) we calculated the probability of FMC being at or below a certain FMC percentile. The conditions used were moderate (−1.5 < SPEI≤−1), severe (−2 < SPEI≤−1.5) and extreme (SPEI≤−2) drought because we were interested in the response to large water deficits (i.e. negative SPEI). These levels are defined by the authors of the drought index (Vicente-Serrano et al. 2010). The FMC percentiles used were 40^th^, 30^th^, 20^th^ and 10^th^ because we were interested in low FMC outcomes.

To assess the reliability of the conditional model in representing the FMC-SPEI relationship, we randomly selected one pixel across each study landscape. We then computed probabilities of FMC on all conditions of SPEI, rather than just the values of interest as earlier. We then normalised the PDF for each SPEI value. We then plotted the normalised probability distribution functions and overlaid the FMC-SPEI observations. If the majority of observations fell within the high probability area, we considered the model reliable (Fang et al. 2019).

We analysed the probability results within landscapes by regression analysis of different layers of conditions of drought and FMC decline with topographical and canopy cover data. Specifically, we used the same FPC data that we used in the non-vegetation masking, and slope, aspect, elevation and topographic wetness index (TWI, all at 30m resolution, Searle et al., 2022). The TWI is a function of catchment area and slope which indicates the potential wetness of a location considering the uphill contributing area (Gallant & Austin 2012). We accessed the topographic rasters through the web portal of the data collection (Table 3, Searle et al., 2022) and reprojected them to the spatial characteristics and coordinate reference system of the model results using the Python packages Rioxarray and Xarray (Hoyer & Hamman 2017, Snow et al. 2024).

## 3. Results

### 3.1 Foliar moisture content variation between and within landscapes

The Pilliga landscape had the lowest maximum FMC of any of the areas, while Severn had a slightly lower minimum than Pilliga, and both landscapes had low means of ∼80% DM (Table 3). In contrast, Kentlyn and Dorrigo had higher means of 126% and 142%, respectively. Overall variation in FMC was greatest at Pilliga with 32% DM (i.e. deviation over space and time), similarly so at Severn with 31%, and lowest at Dorrigo with 9%. Kentlyn had moderate variation of 20% and Murray had moderately high variation of 25%. The results show that landscapes with higher average FMC tend to have lower variation, confirming the landscape scale part of the first hypothesis (H_1_), however Murray defied this trend somewhat with moderately high mean and variation. Through time, all landscapes showed strong and generally synchronised seasonal patterns with winter peaks (midyear, or just after) and summer troughs (Supplementary S1). There was also substantial interannual FMC variation in addition to the seasonal patterns.

FMC patterns were idiosyncratic among the landscapes considered. Spatial patterns of FMC within Kentlyn were distinct, higher means occurred nearer to masked-out drainage lines, where temporal variance was also less (Figure 3). An FMC gradient occurred across a ridgetop in the south-east of the landscape, with higher mean and lower variance surrounding the reservoir in the furthest south-east area. Spatial gradients in FMC at the other four landscapes were less sharp than those at Kentlyn, though there were some areas of Dorrigo with moderately sharp FMC gradients. These gradients differed from gradients at Kentlyn, lower slopes immediately near water courses had lower FMC with wide-ranging variance compared to surrounding higher positions, particularly within the south and eastern ranges. Dorrigo also included areas of relatively high mean FMC with low temporal variance, though variance overall was smallest amongst landscapes. At Pilliga, forests in the south-east, nearby the transition to shrublands, exhibited relatively high FMC with lower temporal variance. At Severn, FMC was higher on average nearer the water body in the north-west and in some distinct patches in central, southern environments, both linear and non-linear in shape. Within Murray, there were strong gradients of FMC mean and variance, with high means occurring with relatively low variance directly around water courses and in wetlands away from the main river channels. The general trend of higher mean and lower temporal variance of FMC near open water confirmed the second part of hypothesis H_1_.

**Figure 3.**
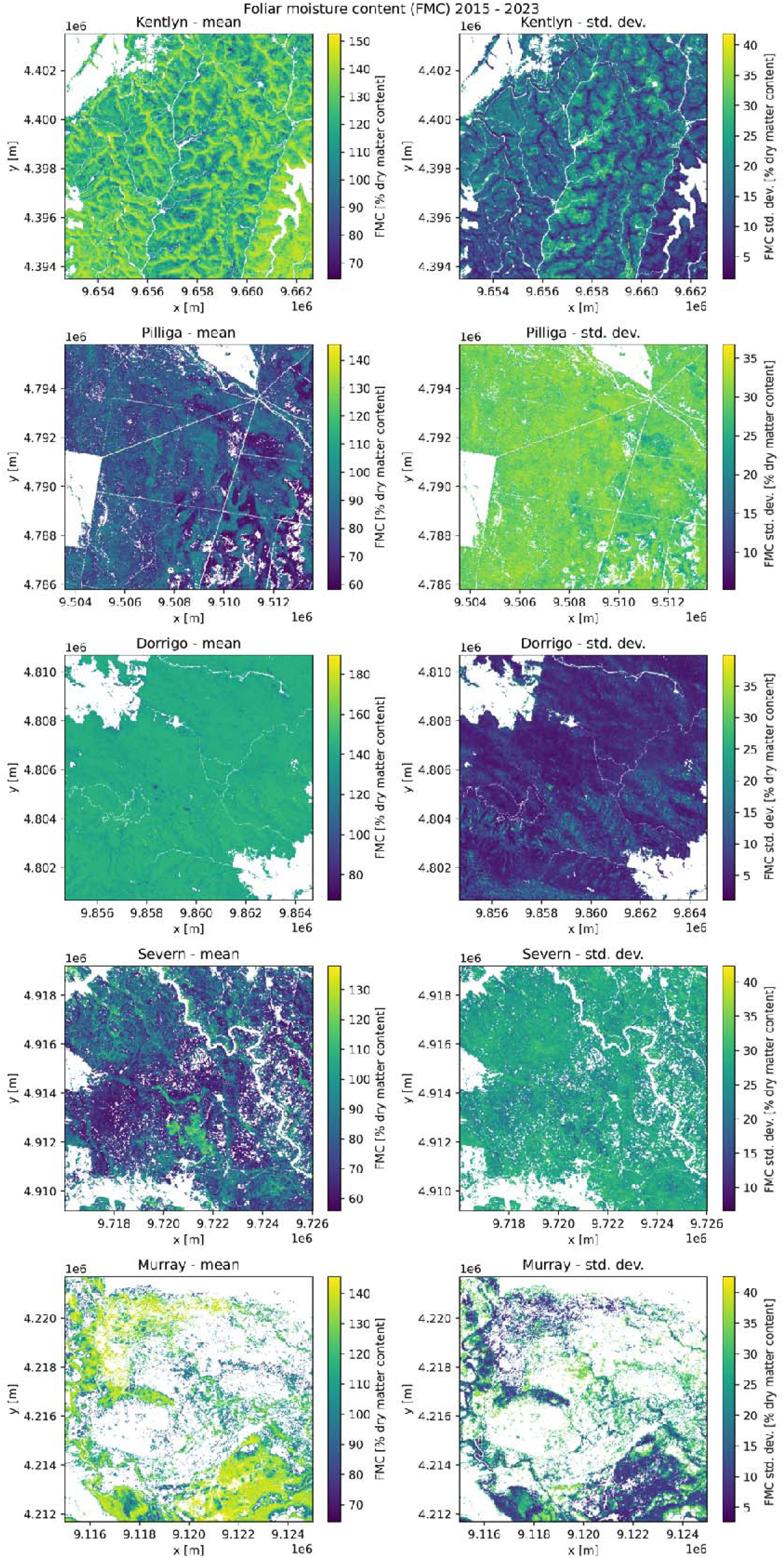
Foliar moisture content mean and standard deviation of forests and woodlands within 5 landscapes in temperate and sub-tropical regions of the state of NSW. Note, each scalebar is different. Distance between axis ticks = 2 km. Landscapes are 10 x 10km in area. Fire scar pixels do not have data covering the full period (2015-2023). Larger version of image available, see link in Data Availability below.

Taking between– and within-landscape changes in FMC together, at Kentlyn the spatial variability approached the temporal variability (Table 4). Whereas Dorrigo had greater spatial variability than temporal variability. At the other landscapes the spatial variability was again less than the temporal variability (∼0.7). These results show that, for FMC, the average variability across each landscape was considerable, relative to the variability through time.

**Table 4.**
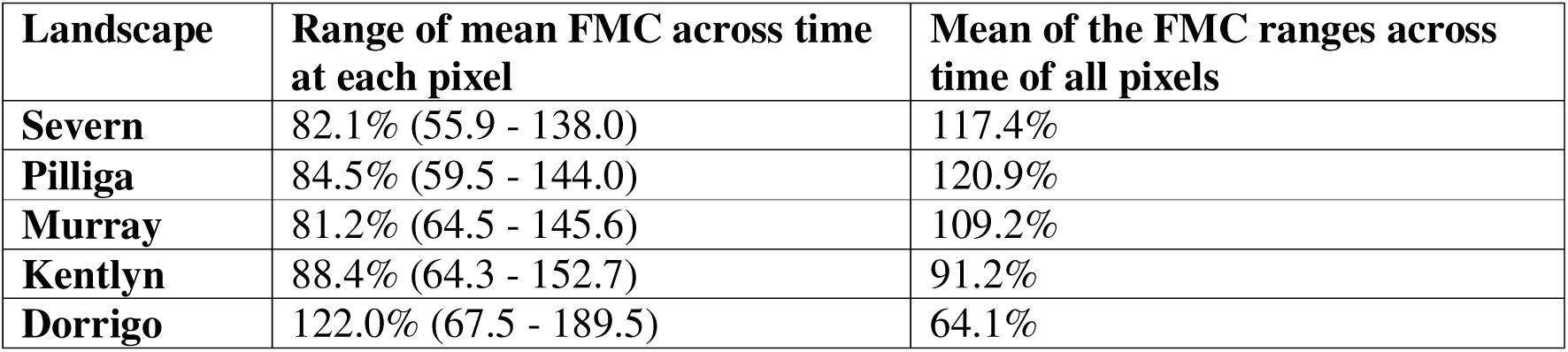
Comparison of spatial and temporal variability within landscapes; range of the mean FMC of each pixel and the mean of the ranges through time of all pixels.

| **Landscape** | **Range of mean FMC across time at each pixel** | **Mean of the FMC ranges across time of all pixels** |
| --- | --- | --- |
| **Severn** | 82.1% (55.9 - 138.0) | 117.4% |
| **Pilliga** | 84.5% (59.5 - 144.0) | 120.9% |
| **Murray** | 81.2% (64.5 - 145.6) | 109.2% |
| **Kentlyn** | 88.4% (64.3 - 152.7) | 91.2% |
| **Dorrigo** | 122.0% (67.5 - 189.5) | 64.1% |

### 3.2 Drought impacts on FMC and associated timescales between and within landscapes

Positive correlations are expected between FMC and climate water deficit because negative SPEI values indicate increased drought. At Pilliga the SPEI effect on FMC was positive, strong and relatively consistent across the landscape, compared to other landscapes (Table 5, Figure 4). Pilliga is the most arid of our sites, thus contradicting our hypothesis (H_2_) that the strongest effect of drought would occur at moderate aridity. At Murray the SPEI effect on FMC was moderately strong, however this varied considerably, across the landscape. At Kentlyn the SPEI effect on FMC was moderate and variable across the landscape. At Severn SPEI had a lesser impact on FMC than other landscapes mentioned but this was variable across the landscape. At Dorrigo the SPEI effect was minimal on average, though there was variation in the landscape. The proportion of forest within landscapes with significant FMC-drought relationships (Figure 5) followed the spatial pattern of positive correlation strength between FMC and SPEI. For instance, at Pilliga 99.6% of pixels were significantly correlated (p < 0.05) whereas 51.8% of pixels at Dorrigo were significantly correlated. These results indicate that the FMC response to drought differs in strength between regions, ranging from little to moderate correlation, and that this differs in strength within landscapes indicating great spatial heterogeneity in forest FMC response to drought.

**Figure 4.**
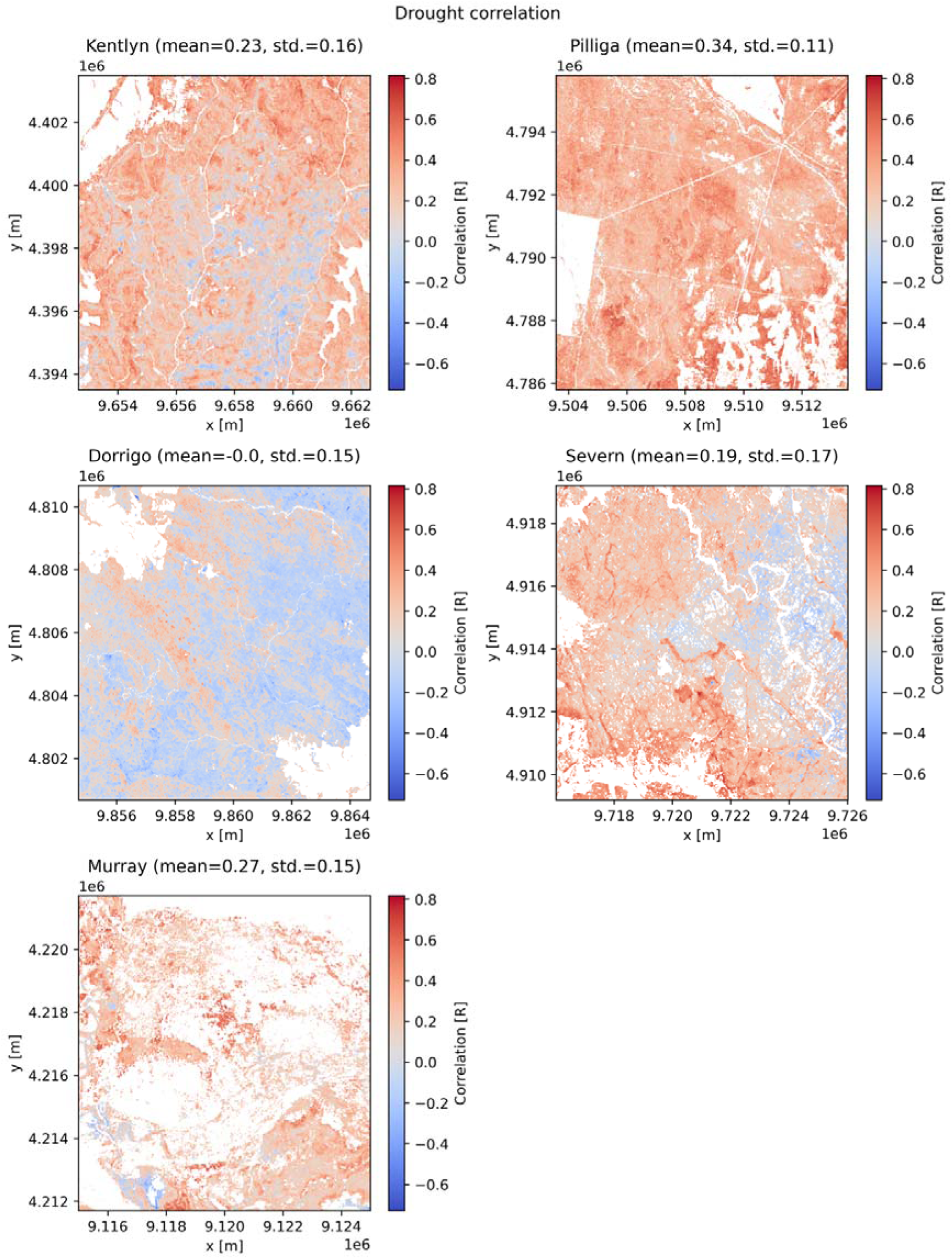
Strongest correlation between FMC and drought (SPEI) at each landscape. Scale is the same in all sub-figures. Distance between axis ticks = 2 km. Landscapes are 10 x 10km in area. Larger version of image available, see link in Data Availability below.

**Figure 5.**
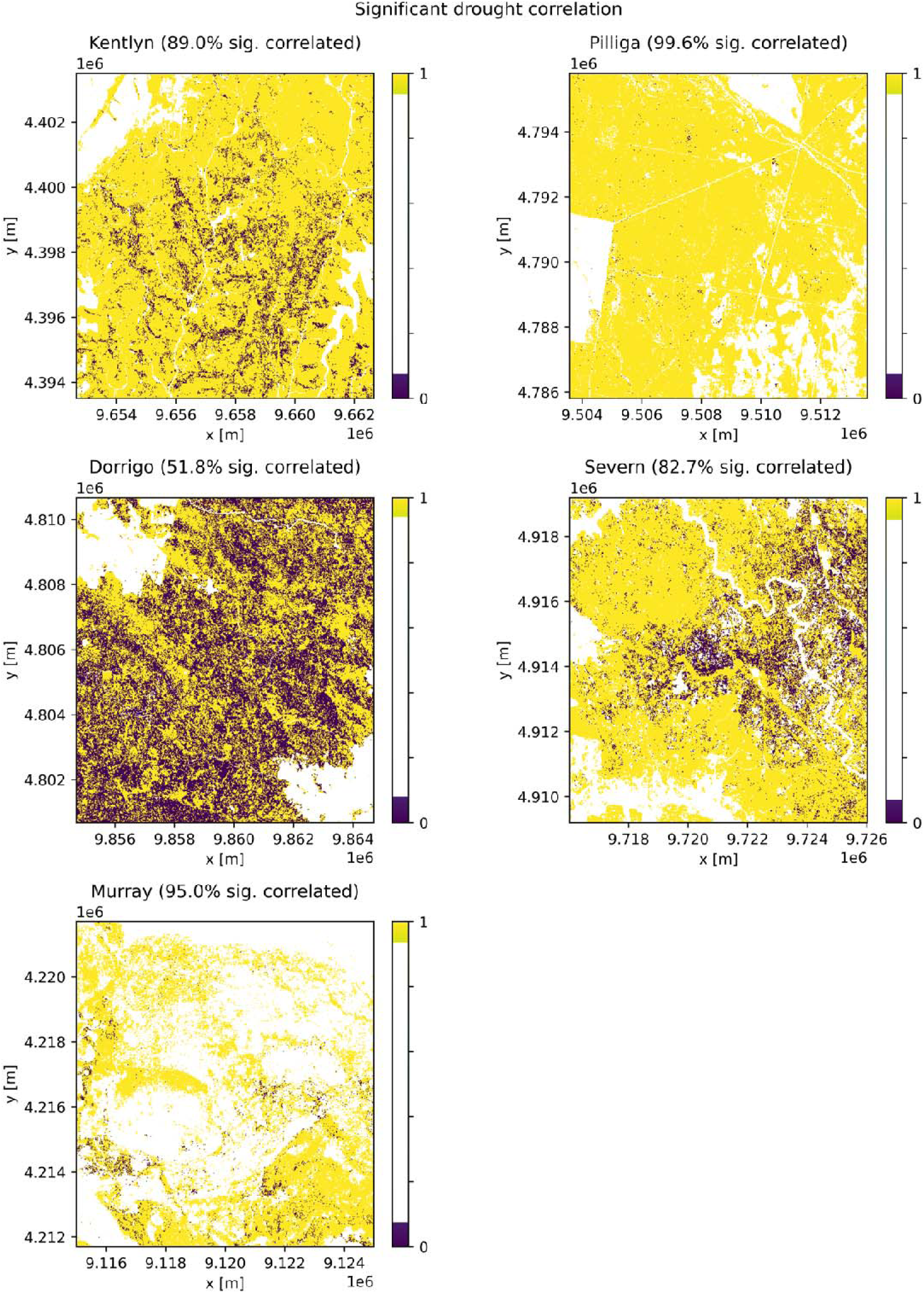
Pixels with (1) or without (0) a significant relationship between FMC and drought (SPEI) of each landscape. Response is Boolean (binary). Distance between axis ticks = 2 km. Landscapes are 10 x 10km in area. Larger version of image available, see link in Data Availability below.

**Table 5.** Landscape means and standard deviations of strongest correlation between FMC and drought (SPEI) and mean and standard deviation of the response time associated with the correlations.

|  | **Drought correlation (Pearson's R)** |  | **SPEI timescale (months)** |  |
| --- | --- | --- | --- | --- |
|  | **Mean** | **Standard dev.** | **Mean** | **Standard dev.** |
| **Pilliga** | 0.34 | 0.11 | 8.38 | 7.41 |
| **Murray** | 0.27 | 0.15 | 20.72 | 16.58 |
| **Kentlyn** | 0.23 | 0.16 | 16.73 | 10.67 |
| **Severn** | 0.19 | 0.17 | 12.90 | 13.85 |
| **Dorrigo** | -0.00 | 0.15 | 18.30 | 15.31 |

At Pilliga, the FMC response time to changes in the climatic water balance (i.e. SPEI timescale) was rapid, on average (Figure 6). The response time was slower at Murray on average, though this varied across the landscape. The response time was moderately slow and relatively consistent across Kentlyn. FMC at Severn responded strongest to moderately long SPEI timescales on average, though this is also variable across the landscape. At Dorrigo responses in FMC to SPEI generally took the longest time to occur, among sites. These results suggest that the strongest responses to climatic water changes occur over different timescales in different parts of forests, with the standard deviation approaching or greater than the mean response time for all landscapes.

**Figure 6.**
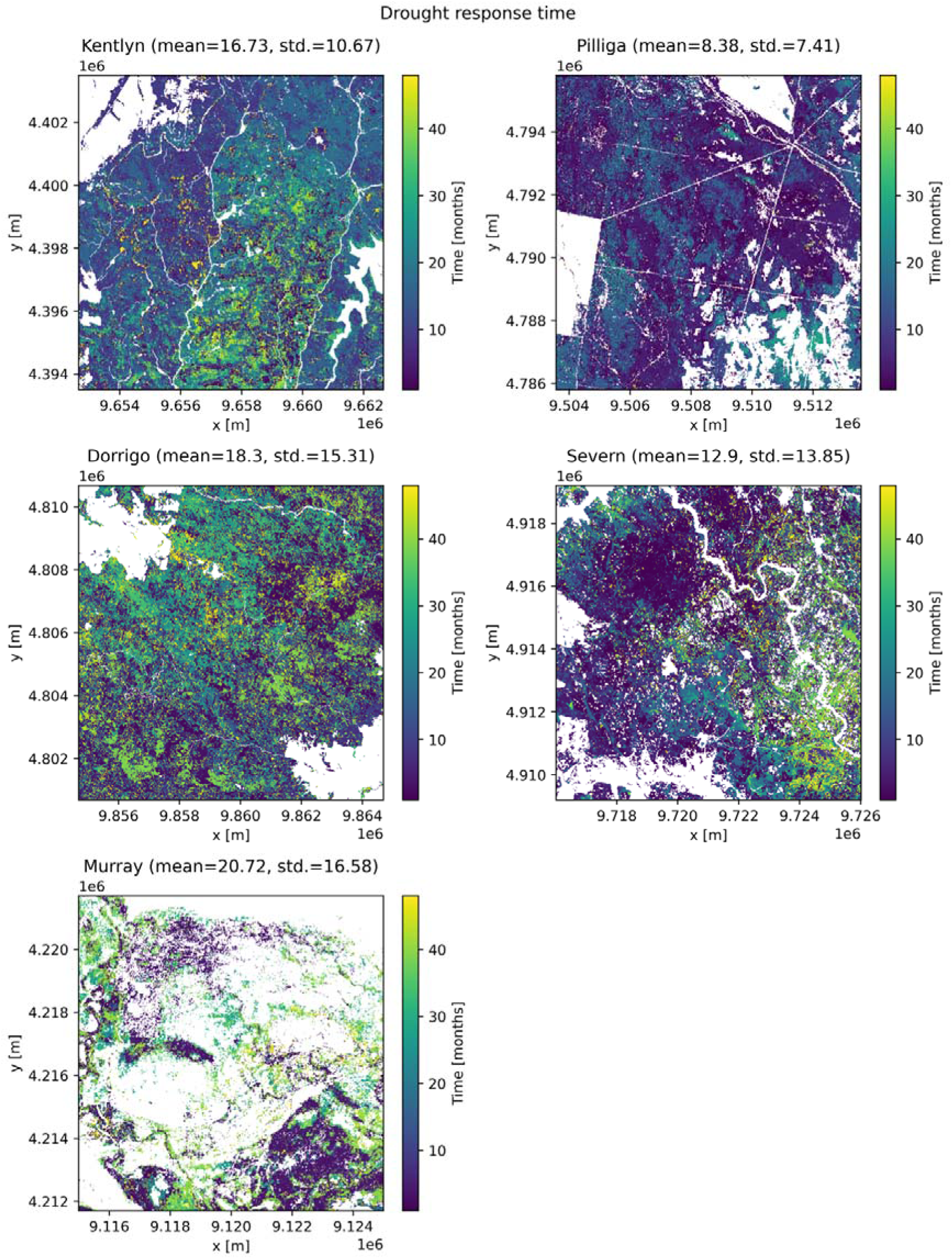
SPEI time scale associated with the strongest correlation with FMC. Scale bar is the same in all sub-figures. Distance between axis ticks = 2 km. Landscapes are 10 x 10km in area. Larger version of image available, see link in Data Availability below.

Within Pilliga, FMC decline with declining water balance was strongest in the south-east in forest surrounding the areas dominated by shrublands (masked, Figure 4). Similarly, so in patches in the south-west and centre of the landscape. These patches of strongest SPEI effect on FMC tended to have a short to medium response time (6-15 months) whereas the patches near the shrublands had longer response times (20-30 months, Figure 6).

Within Murray the water balance effect on FMC was weakest (or negative) in patches of riparian forest very close to water courses, however there were also areas of relatively strong positive correlation at some places along riverbanks (Figure 4). Moreover, the highest positive correlations of FMC with SPEI tended not to be in forest fronting the main river channels. The response times of the forests most influenced by the climatic water balance were fast (1 or 2 months), in large areas of wetlands away from the main river channels. Along the main river channels there was a range of SPEI response times (up to 40 months) (Figure 6).

Within Kentlyn the SPEI effect on FMC was positively strongest in forests of the north, east, and west, apparently influenced by the topography of river valleys (Figure 4). In general, these higher positive correlation forests had shorter response times, while the central area of weak positive and negative correlations had long drought response times (Figure 6). Within Severn strongest positive FMC-SPEI relationships occurred in several small patches or strips amongst rocky outcrops, also in areas bordering cleared vegetation in the south (Figure 4). The SPEI time scale was variable for these forests showing high SPEI correlations, with some of the forests in gullies showing long timescales for strong drought effects (Figure 6). Within Dorrigo the relationship of FMC and drought was positively strongest in the central to north-west area (Figure 4), however most of the landscape was negatively correlated or not at all. There were localised areas of high positive correlation in the other parts of the landscape, apparently varying with topographies.

The landscape average relationships of FMC with different SPEI timescales, of up to 48 months, are shown in Figure 7 (study period = 92 months). Though FMC at Pilliga did respond most strongly to SPEI of a relatively short timescale (∼8 months), only slightly weaker relationships occurred for SPEI of 15-20 months, beyond this length of time there was a decline in the relationships. A similar pattern in FMC-SPEI correlation with SPEI timescale occurred at Severn. At Kentlyn, FMC did not relate strongly to SPEI of short timescales (1 or 2 months), but this changed quickly to a strong relationship over an 8-month timescale. The strength of relationships at Murray also increased quickly over the shortest timescales but with lesser magnitude than Kentlyn. Dorrigo also tended to demonstrate an increases in the strength of correlations, though gradually, up to a relatively long SPEI timescale (25 months). Dorrigo also showed two peaks in FMC-drought correlation for two widely differing drought response times (∼ 5 months and > 2 years).

**Figure 7.**
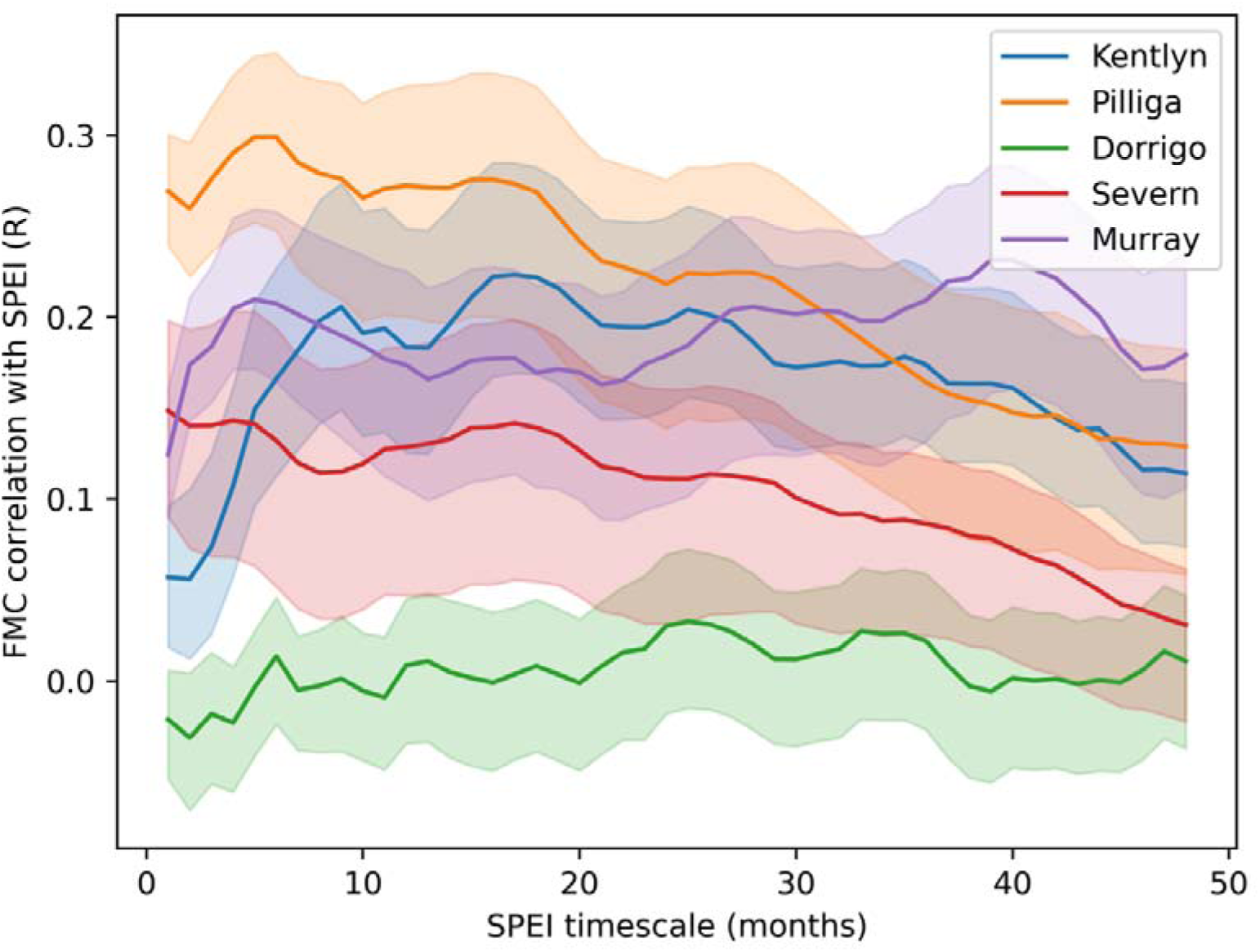
The relationship of FMC to drought (SPEI) for differing timescales of SPEI. FMC correlation is the Pearson’s correlation coefficient (i.e. Pearson’s R) averaged for each landscape. Error bars are half a standard deviation. SPEI timescales are in monthly increments from 1 to 48 months.

### 3.3 Probabilities of FMC decline with drought between and within landscapes

The probability of 10^th^ percentile FMC during extreme drought was lowest at Dorrigo and Kentlyn (both mean *P=*0.09), and highest at Pilliga (mean *P=*0.18, Figure 8). These results are for the SPEI timescale of greatest FMC-SPEI correlation (Figure 6), and confirmed our hypothesis that low FMC levels are generally rare (H_3_). This result indicates that during extreme climate water deficit FMC at Pilliga is twice as likely to decline to the lowest levels compared to Dorrigo and Kentlyn. The masking of FMC data within two years after fire altered the probability results greatly within landscapes, therefore we masked out fire scars completely in these results (Figure 8). The variation in probability of FMC decline was largest at Pilliga and smallest at Kentlyn (std. dev. *P=*0.08 and 0.02 respectively). This shows that despite the high likelihood of FMC decline at Pilliga there is great variation within the landscape, including areas where the lowest FMC levels are less likely to occur in extreme drought, compared to Kentlyn where the chances of decline are more spatially consistent. Though even at Kentlyn and certainly at Pillliga there was high dispersion in the probability of decline (CV = 22% and CV = 44%, Figure 8), confirming our hypothesis that variation across fine environmental gradients is high (H_4_). Severn had higher spatial variation in probability of decline alongside a higher average, like Pilliga. We found the actual values of the 10^th^ percentile FMC were lowest at Pilliga, then higher at Severn, then Murray, followed by Kentlyn and Dorrigo, on average (Supplementary S2). This result shows that actual quantities of FMC in forests are very different across landscapes for the same FMC percentiles. The probabilities of FMC declining to percentiles higher the 10^th^ (i.e. 20^th^, 30^th^ or 40^th^), were generally higher for any level of drought (data not shown).

**Figure 8.**
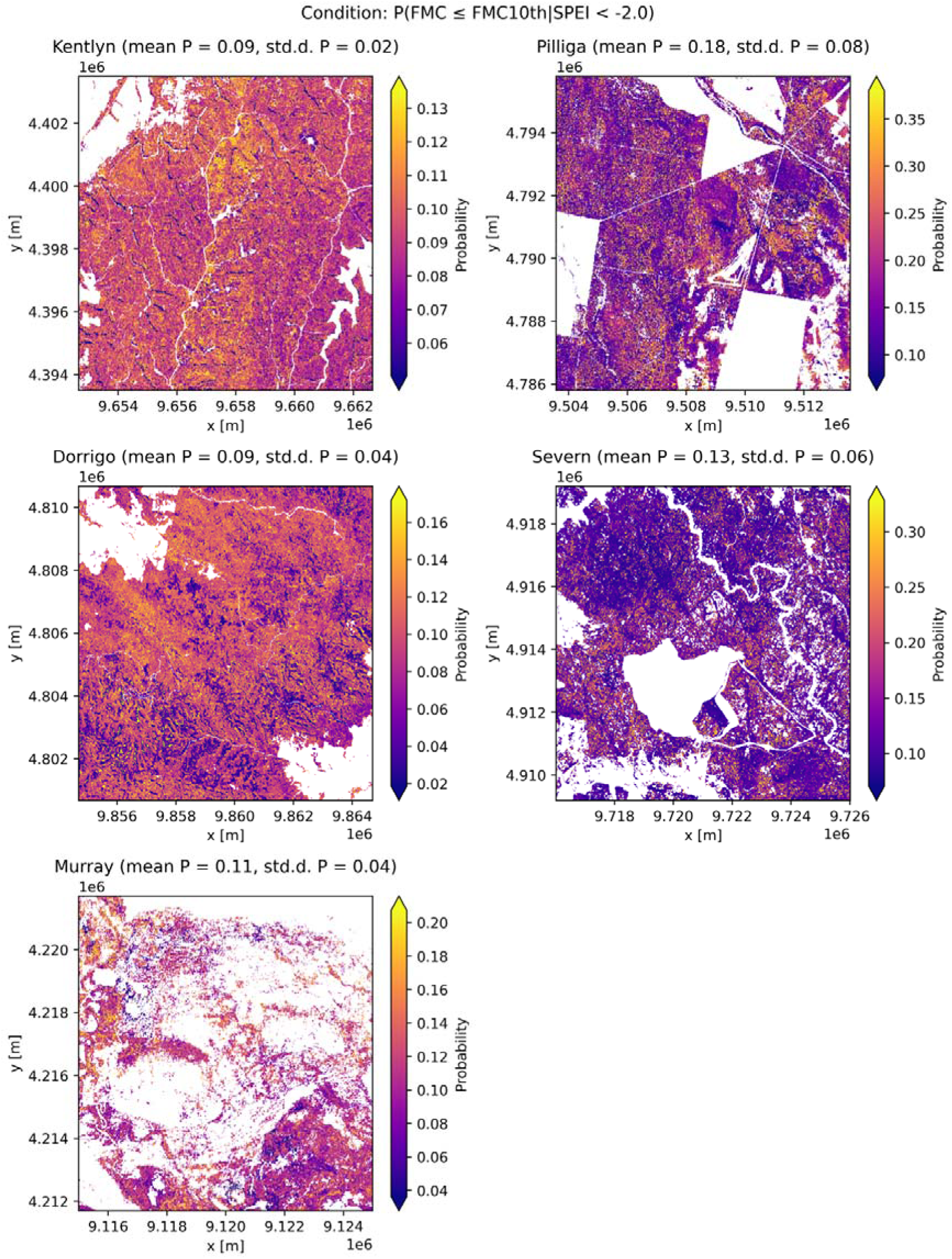
Probabilities of great foliar moisture loss (i.e. ≤ 10^th^ percentile) under extreme drought, across each landscape. The scale bars are different for each landscape. Some areas have been masked which were mapped as fire scars for fires that occurred during the study period (particularly Pilliga and Severn). Distance between axis ticks = 2 km. Landscapes are 10 x 10km in area. Larger version of image available, see link in Data Availability below.

Considering again declines to the 10^th^ percentile FMC, areas of low probability of FMC decline within Kentlyn occur as discreet strips of forest in river valleys or creek lines and south facing slopes particularly in the west and north (Figure 8). At Pilliga areas of low probability of decline occurred along the borders of the shrublands in the south-east and in patches of dense forest in several locations, sometimes along creek lines. The areas in between ephemeral creek lines in the central-north-east part of the landscape hold two of the patches of forest with low chance of FMC decline with extreme drought. At Severn, the south-west facing slopes of Severn River have relatively low probability of decline, also in the area surrounding a reservoir and nearby creeks. Also, in several small valleys between rocky outcrops FMC is likely to be relatively maintained in extreme drought compared to other parts of the landscape. At Dorrigo several patches of forest in the centre and north-east have relatively low probability of FMC decline, also in many valleys in the southern section of the landscape. In these valleys the forest of lower chance of FMC decline in extreme drought do not necessarily occur on creeks or rivers but back away from them in many cases. Forest of highest probability FMC decline at Murray occurred further away from watercourses however there was a range of probabilities along the main channels, while the wetland areas in the north and south of the landscape tended to have lower chance of FMC decline to 10^th^ percentile in extreme drought.

We found very little correlation between the probability of FMC decline to low levels and topographic variables (elevation, slope, aspect, TWI, data not shown), the strongest being a positive relationship between elevation and probability of FMC decline at Dorrigo (R = 0.13, r^2^ = 0.02, *P<*0.05). This result suggests lower elevation forests have FMC which persists in extreme drought relative to higher elevations, in landscapes with a wide range of elevations (i.e. ∼50 to ∼1000m). The next strongest relationship was a negative relationship between slope and probability of decline at both Murray and Severn (R = –0.08, r2 = 0.01, *P<*0.05 and R = –0.07, r2 = 0.0, *P<*0.05 respectively). These results suggest that steeper slopes allow to a small degree for FMC to persist in extreme drought, at some landscapes. However, these results show that chance of great FMC decline in extreme drought is highly variable within landscapes. We found similarly weak relationships between topographic variables and the mean of FMC across time (data not shown). This further emphasises the variability in FMC within landscapes.

Within Murray, Kentlyn and, particularly, Dorrigo, as mean FMC increased the chance of FMC decline with extreme drought, lessened (Figure 9). However, at Pilliga and Severn there were two distinct clusters within this mean FMC-probability space. One cluster was somewhat like the other landscapes with flat or negative slopes, generally having slightly lower probability of FMC decline in drought with increasing average FMC. The other cluster tended to have an increasing chance of FMC decline in drought with increasing mean FMC and did not have more than moderate mean FMC levels. Pixels of this latter cluster tended to occur around the edges of relatively contiguous patches of the former cluster.

**Figure 9.**
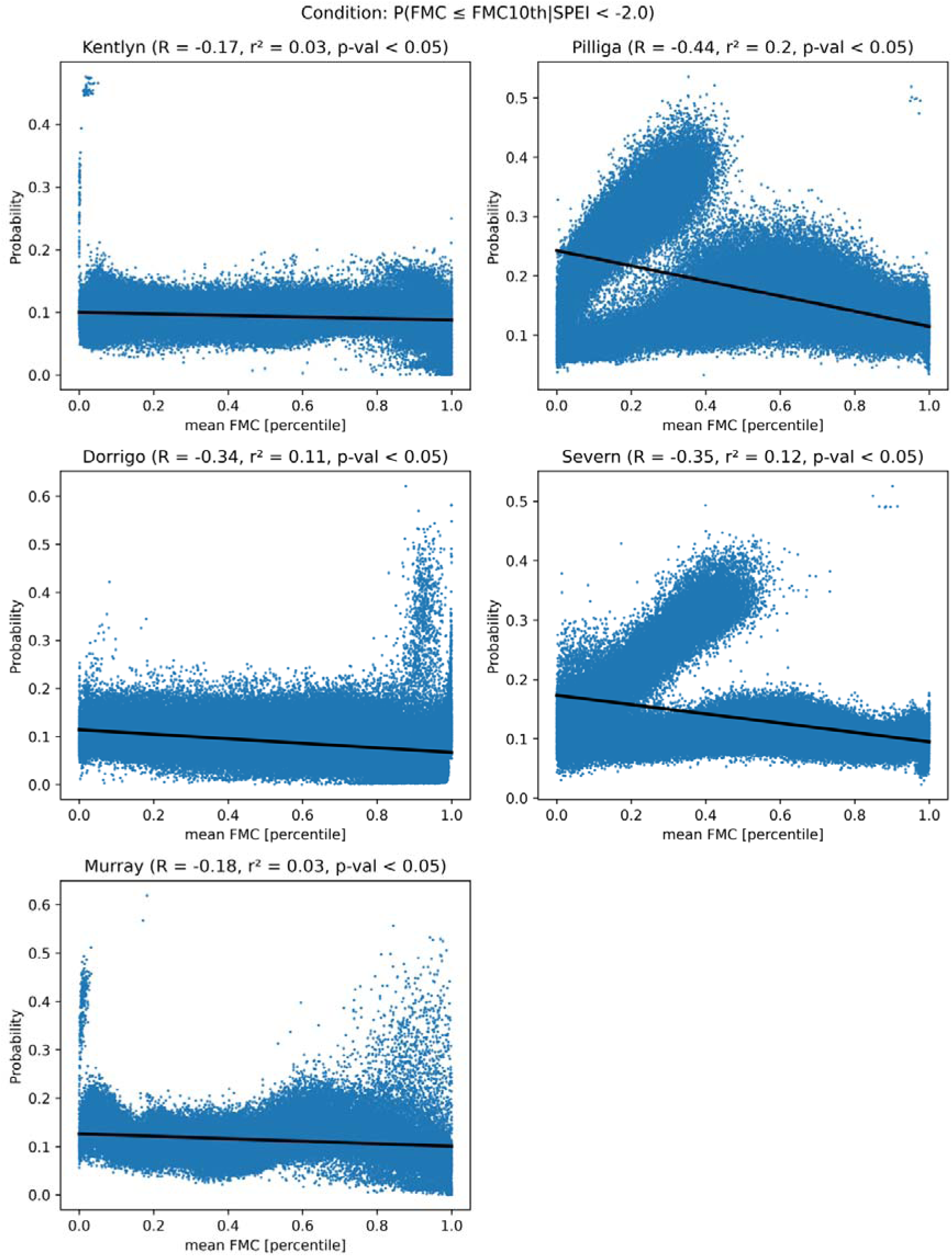
Linear regression plots of probability of great foliar moisture loss (i.e. ≤ 10^th^ percentile) under extreme drought against mean FMC at landscapes.

We found that sites in Kentlyn which have a low probability of drying out are consistently those with higher FPC (R=-0.32, r^2^ = 0.10, *P<*0.05, Supplementary S3), and this, strongest, effect is for moderate FMC declines (i.e. 30^th^ percentile). There was also a relationship between FPC and probability of decline at Dorrigo, the strongest being for the lowest FMC and moderate drought conditions (R=-0.26, r^2^ = 0.07, *P<*0.05). Other landscapes did not have a notable relationship between foliage cover and chances of FMC decline.

## 4. Discussion

This study utilised an extensive FMC dataset combined with meteorological data to understand relationships with drought and identify forests and woodlands of persistent FMC, indicating resistance to drought.

### 4.1 FMC across and within landscapes

There was great spatial variation in FMC of forest and woodland landscapes. The variation within landscapes was generally great relative to the range through time. This relatively high spatial variation occurred in mesic and arid landscapes. More broadly we captured FMC differences between landscapes which align with average rainfall, as was shown by Nolan et al. (2022) for maximum FMC. We also found declining FMC variation with increasing rainfall (H_1_), except in the landscape Murray, which had the lowest rainfall but relatively high average FMC, due to the riverine context which is influenced greatly by varying river flows (Selwood et al. 2019). We found fine-scale changes in FMC which can be explained by environmental gradients, within all landscapes.

Within Kentlyn, forest FMC changed quickly over short distances close to water courses due to increased soil moisture on lower slopes, which has been shown to influence plant water relations via depth to groundwater (Zolfaghar et al. 2015), and due to the protection from wind and radiation created by the incised gorges of the sandstone-shale geology (Sluiter et al. 2002). Also, FMC was more stable in some of the south-east valleys due to aspects that receive ocean-associated moisture and rainfall, and less radiation. A lack of these characteristics, i.e. exposure, can contribute to tree desiccation and dieback (Fitzgerald et al. 2023). This sheltering effect also explains the relatively high FMC, sclerophyll, forests of the south-east facing valleys within Dorrigo. Across this site there was an overrepresentation of FMC values of ∼140%, and not many values exceeding this level, however this is within observed FMC distributions of rainforest species which dominate the landscape (Krix & Murray 2018). This congruence is despite the FMC model not being calibrated or validated for such species. Within Severn, distinct high mean FMC areas in the lower centre, which appear to be in the narrow areas between rocky outcrops, as seen in satellite imagery, are due to deeper and more moist soils of these drainage lines allowing higher plant FMC. This effect is reflected in the occurrence, there, of relatively dense woodland communities compared to surrounding environments (Hunter 2015). The high-mean, low-variance FMC patches bordering shrublands at Pilliga are likely groundwater accessing woodlands associated with the Pilliga outwash ecosystems. These woodlands grow in sandy soils at the edge of colluvial deposits where water moving from upslope comes toward the surface in these deposits or accumulates across the surface to these lower-slope environments, facilitating higher, stable FMC compared to the drier communities, upslope (Ickowicz et al. 2018, Benson et al. 2010). High-mean, low-variance FMC also occurred at Murray, in wetlands and along water courses, which is likely due to trees’ access to the water table and specifically to the river channels themselves. In these environments *Eucalyptus camaldulensis* forests grow and are dependent on the river flows and associated soil moisture (i.e. phreatophytic, Doody et al., 2014).

### 4.2 Drought impacts on FMC and associated timescales

The high spatial variability in strength of SPEI effect on FMC within sites is likely due to species and environmental differences which regulate trees uptake and loss of water (Nolan et al. 2020), some of these differences within sites were mentioned previously in relation to FMC patterns. Given the importance of water for tree growth and productivity we expected strong FMC relationships with variation in the water balance, particularly in moderately arid landscapes (H_2_) where plants experience lower soil moisture and higher atmospheric demand (Nolan et al. 2022). We did find this to be the case, however the most arid sites had the strongest correlations between SPEI and FMC on average. This may be due to the strong water deficits of arid landscapes having a more pronounced influence on canopy response than species influences, as expected.

Within Pilliga, some strongly SPEI affected forest patches had differing SPEI timescales, likely due to upper-slope drylands with sandy soils declining in moisture (Ickowicz et al. 2018) faster (i.e. 6 to 15 months) than the lower slope forests near the washout areas, which took longer to be strongly affected by SPEI changes (i.e. 20-30 months). These SPEI response timescales must also include the soil and plant influences and not only climatic moisture variations. Pilliga contains drought adapted tree species (e.g. Ironbarks, *Eucalyptus sect. Adnataria*), amongst others (e.g. *Callitris sp.*), that regulate their water balance efficiently compared to more mesic adapted species (Silva et al., 2016, *unpublished manuscript*). However, the average timescale of strongest SPEI effects at Pilliga was relatively short, 8 months. This response time may reflect arid adapted species having less moisture to lose during drought because of a tight regulation of water in the leaves, in general, leading to a relatively quick shift to a droughted state compared to mesic species and forests. For instance, diel FMC declines are smaller in species of lower average FMC and drier climates of origin, including during droughts and heatwaves (*unpublished manuscript*). This smaller, faster type of FMC response may have also occurred in the arid Severn landscape that had low FMC and the second shortest SPEI timescale, on average.

Other tree adaptations may contribute to spatial differences in the SPEI effects on FMC, alongside declines in leaf FMC, rapid or otherwise. These processes include reduced leaf area by increased excision and slowed growth of leaves during drought (Pritzkow et al. 2020). Such response to negative SPEI likely contributed to the weaker SPEI-FMC relationships observed within some areas of Pilliga, Severn, Murray and Kentlyn. The loss of leaf area will reduce canopy level FMC, but in relatively low FPC sites (Griebel et al. 2023) more of the understorey signal is introduced into the reflectance (Pereira et al. 2004), influencing the FMC estimates (Kotzur et al. 2025) and SPEI response. Previous studies have been limited in detecting leaf loss as a drought resistance strategy due to the covariance of canopy moisture with leaf area (Brodrick, Anderegg and Asner, 2019). However, the covariance is accounted for in the RTM model of FMC used here, as canopy simulations include a range of leaf area for each FMC level, any of which can be retrieved for the same pixel across time (Kotzur et al. 2025). Though there is a lower limit to leaf area at which canopy reflectance will be lost. The effectiveness across time of this leaf area retrieval is yet to be explicitly tested.

At the mesic end of the climatic gradient, in Dorrigo, the common rainforest patches had high FMC, limited SPEI response, and long average, though extremely spatially variable, SPEI timescales. The long SPEI timescales reflect that rainforests are resistant to droughts, mainly because they are adapted to environments which rarely experience droughts (Tng et al. 2018). Some of the patches of forest that did have a significant positive relationship with SPEI, over relatively long timescales, were on western facing aspects which are more exposed. This agrees with recent observations of rainforests drying to the point of rare flammability in contemporary extreme droughts (Collins et al. 2021).

Murray tended to respond positively to SPEI over moderately long timescales (12 months, i.e. 12 months longer than Pilliga), due to the riverine environment preserving water supply longer into drought than more arid, upslope landscapes. Also, Murray is dominated by *E. camaldulensis* trees that have deep roots able to access the river and water table (Selwood et al. 2019). However recent reduction in river flows by humans could be expected to start reducing FMC at less negative SPEI levels (Doody et al. 2014), hence strengthening the SPEI-FMC relationship. Though this may occur over longer droughts than was studied here.

In this study, the FMC-SPEI relationships included non-drought periods (i.e. positive SPEI) that may obscure, strictly, drought responses (i.e. negative SPEI) if the FMC responses differ greatly between positive and negative SPEI values. For example, tree water potential can recover within 3 weeks of cessation of severe drought (Gauthey et al. 2022), though leaf loss may alter this. Beyond the positive SPEI levels necessary for this recovery, FMC may increase little, thereby lessening the strength of the FMC relationship overall, regardless of a strong negative SPEI response (i.e. drought).

The relationships of increasing forest FMC in drier conditions (i.e. negative correlations, Figure 4) were predominantly in the most mesic landscape, Dorrigo, which had the highest and most stable FMC overall. Many of these negative SPEI-FMC relationships were not significant and the strengths were not as great as for positive relationships (Figure 5, Figure 6). Negative SPEI-FMC relationships in other landscapes were associated with shrublands (Severn) or heathland and open woodlands (Kentlyn), both being on rocky outcrops which may convolute FMC-SPEI relationships, particularly because shallow soils tend to facilitate low leaf area (Fisher et al. 2016), hence a weaker tree reflectance signal.

### 4.3 Probabilities of FMC decline with drought

Overall, the probabilities of tree canopy FMC declining to the 10^th^ percentile or lower during drought, were low (average 9 to 18% amongst sites), owing to the inherent rarity of extreme droughts (H_3_), however there was substantial variation within sites (H_4_). The probabilities of decline increased, on average, from mesic to arid sites, except at the climatically arid, riverine landscape (Murray) which had lower probabilities than might be expected, for reasons mentioned previously. In other landscapes, those which were more arid had larger variation in probabilities of decline and lower absolute FMC levels than mesic landscapes (Supplementary S2). These results show that trees of relatively arid landscapes have greater adaptations for climatic drought to exploit relatively small water resources of various microenvironments, namely through more extensive tree hydraulic systems (i.e. roots and vessels) and stricter control of plant water potential (e.g. stomatal regulation, Choat et al., 2018). This capacity in forests and woodlands of arid landscapes allows greater spatial variation in resistance to drought than mesic landscapes (Selwood et al. 2019). High variation in probability of FMC declines also occurred in the Murray landscape. Murray is climatically dry compared to other landscapes but is comprised of riverine ecosystems. Therefore, the high variation of FMC decline may be influenced by controlled unseasonal river flows, the extents of flooding (Treby & Carnell 2023), and the proximity of trees to river channels where climate resilience tends to be higher (Selwood et al. 2019). High relief landscapes showed distinct, more linear, patterns in forests with drought persistent FMC, these were associated with valleys and aspects therein. Whereas in low relief landscapes, forests with low probability of FMC decline were in disparately shaped patches (Figure 8). At the most arid, low relief sites (Pilliga, Severn) the forest and woodlands where FMC was retained during drought were found both near and far from creeks and rivers. The reasons for these landscape patterns are likely due to the soil and slope effects mentioned earlier. Our results broadly agree with (Brodrick, Anderegg and Asner, 2019) who showed forests in the state of California (USA) to have “significant spatial heterogeneity in drought resistance” as quantified by a meteorological index and remotely sensed canopy water content. As we did, they found greater resistance to drought, on average, in mesic than in arid forests.

### 4.4 Ecological implications

Our results are important in understanding distributions of arboreal herbivores facing dehydration during drought (Briscoe et al. 2016), events which are expected to increase in frequency due to climate change, across the study region (Zhongming et al. 2021). Despite having the greatest average chance of FMC decline in drought, Pilliga had the greatest spatial variability in this chance, indicating that local refugia are present. These refugia were in larger patches (∼10 to ∼70 ha) and smaller patches (< 2 ha) interspersed in pixels of high probability of FMC decline throughout the landscape, revealing the locations of a crucial environmental resource during drought. For folivores, these forest patches mean greater water availability in leaf diets (Briscoe et al. 2016), assuming that species are adapted to the local (Ellis et al., 1995), absolute FMC levels, which decreased with landscape aridity. Severn was the landscape next most spatially variable in chance of FMC decline with drought, after Pilliga, and both these landscapes were the most affected by shorter SPEI timescales, indicating that forests of faster FMC responses to drought have more patches of local refugia, relative to other, more mesic landscapes which are less responsive and more homogenous in chance of FMC declines with drought. Animals may be more selective for forage moisture (Melzer 1994) due to these refugial forests, potentially changing the number or composition of trees used to include those of the refugial patches, as is done for other important nutrients (Moore & Foley 2005). These preferences have been observed in koalas, they chose trees near water courses during drought, expanding into surrounding habitat trees in wetter conditions (Smith et al. 2013). Such preference changes in populations may increase density, competition between animals for resourceful habitat, and increase browsing pressure on trees themselves (Whisson et al. 2016). For insects, the patches of forests which have lesser chance of FMC decline will be of increased habitat quality because of higher leaf water content and potentially greater leaf area as drought progresses compared to surrounding forests. Particularly so for sap-sucking insects which require turgid leaves to access nutrients (e.g. psyllids, Gely, Laurance and Stork, 2020a). These higher quality patches could result in high population growth and resultant defoliation (Angel et al. 2008), which would then reduce canopy level FMC. Defoliation also affects the reflectance signal of the tree canopy (Masaitis et al. 2013), likely causing erroneous FMC estimates (Kotzur et al. 2025). However, at lower levels of drought, tree chemical defences can increase, potentially lessening the quality of this refugia for insects (Gely et al. 2020).

The drought resistant forests and forest patches revealed in this study are important for understanding forest fire risk which is strongly affected by the dryness of forests, and forest biomass (Nolan et al. 2016). This is particularly relevant in the wetter landscapes where biomass is higher, but the spread of fire may be impeded by forest patches that are too moist to be flammable (Krix & Murray 2018). Whereas, within more arid landscapes, lower absolute levels of FMC allow fire spread in dry conditions, as reflected in the large extents of burned area from bushfires in recent years (Boer et al. 2020). Indeed, Rao et al. (2022) used ‘plant water sensitivity’, a similar metric to the probability-decline results presented here, to show that with an increase in atmospheric aridity (VPD), burned area increases more in vegetation of higher plant water sensitivity (i.e. greater chance of FMC decline with drought).

The drought sensitive forests identified here, i.e. higher chance of great FMC decline in extreme drought, include those experiencing canopy dieback, which is also important for koala habitat quality. Dieback may occur when the tension of water within the xylem of trees increases, due to drought driven reductions in soil moisture often coupled with increases in air temperature and evaporative demand, to the point where air bubbles form *en masse* blocking water from passing up to the canopy and damaging this hydraulic system (Choat et al. 2018). This phenomenon is evident in the satellite-based FMC response, such as for documented forest dieback during the 2019 drought in New South Wales, Australia (Losso et al. 2022). The recovery signal from dieback appears to be hastened, possible by a prominent understorey signal after canopy desiccation and droughts end. Dieback entails a loss of moisture available to folivores, along with most other forage nutrients, which may be fatal by malnutrition and dehydration if enough trees within habitats are affected (Whisson et al. 2016). Repeated dieback due to frequent extreme droughts may also be the driver by which the distribution of koala food trees is progressively altered, distributions have been predicted to change mainly by contraction southward and eastward with some minor expansion outside the current range (Adams-Hosking et al. 2012). The FMC data cube contains the potential to analyse dieback and changes in the distribution of species’ habitat.

### 4.5 Study caveats

We have identified patterns in forest FMC and the probability of its decline with drought however our results may be biased due to the influence of understorey vegetation on canopy signals, though we controlled some of this error with FPC masking of lower canopy density pixels (Kotzur et al. 2025). The bias was apparent in the landscapes of lower canopy density (i.e. Pilliga and Severn), specifically in the relationship between mean FMC with probability of substantial FMC decline in extreme drought, within these landscapes (Figure 9). There was a distinct group which had low to moderate mean FMC and increased probability of decline with mean FMC. These pixels tended to occur at the edge of higher mean FMC patches and had low to moderate foliage cover. This may result from a signal of understorey shrubs and grasses that tend to have lower FMC than forests (Yebra et al. 2024). Although the accuracy of the FMC estimates in these vegetation types is not known (Kotzur et al. 2025). Shrublands and grasslands have less extensive root networks than trees, thereby exploiting less soil volume for water in drying soils (Schenk & Jackson 2002), though root function also plays a role in this process (Nippert & Holdo 2015). Whereas the, presumably, exclusively tree FMC signal showed more persistent FMC during drought in pixels of higher average FMC. This trend being due to the decoupling of trees, unlike shrubs and grasses, from shallow soil moisture (Brown et al. 2022), facilitating resistance to drought stress (Zolfaghar et al. 2015). This problem of understorey signals has been documented in remote sensing studies; it is inherent to optical sensors and limits applications in ecosystems with sparse canopies (Nolan et al. 2016, Eriksson et al. 2006). The problem may be reduced by increased spatial resolution of reflectance data to mask sparse canopies or the gaps therein (e.g. 3 m resolution with LiDAR based mask, Dixon et al. 2023). Additionally, this study did include some vegetation types other than forest or woodlands, in some areas of Severn and Kentlyn, due to inaccuracies of the forest height and FPC layers. Furthermore, rainforest vegetation types were included in the study as they make up most of the Dorrigo landscape. There, small parts of the rainforest showed unexpectedly low FMC, often on lower slopes, compared with surrounding eucalypt forests (Krix & Murray 2018), which may be caused by the model not having been parameterised in rainforest.

### 4.6 Future work

Our modelling approach is suited to forecasting FMC. Forecasts of precipitation and potential evapotranspiration from climate models could be computed into a drought index, which would facilitate the probabilistic estimation of future FMC decline based on the dependence within each pixel, modelled here, or the dependence modelled including subsequent FMC observations. This assumes that the dependence structure between FMC and SPEI will not change in future, however hydro-physiological limits of forests are being tested by drought in many biomes (Choat et al. 2018). Copula approaches have been used in combination with forecasts from climate models. For example, Goswami et al. (2018) predicted likelihood of joint, extreme precipitation events over North Sikkim Himalaya. However, the prediction was based on the comparison of likelihood of future joint-events and that of past joint-events, rather than using the dependence of the past to predict the events (or probabilities) of the future. Regardless, the potential for forecasting FMC exists, and would be important information for environmental managers. For instance, fire extent is largely driven by foliar moisture conditions, as mentioned earlier, which can change rapidly in time (Boer et al. 2017, Nolan et al. 2016). The forecasting would also be useful in biodiversity conservation, as is the case in predicting heat stress in flying fox for planning management interventions (*Pteropus spp.,* Ratnayake et al., 2019). Such forecasts of FMC and the dataset analysed here can be used in further analyses to understand FMC gradients. Particularly, the relationship of hydrological drought and FMC could be analysed with more directly associated variables such as soil moisture (e.g. Zolfaghar et al., 2015; Vinodkumar et al., 2021) and vapour pressure deficit (Griebel et al. 2023), to provide greater explanation of variation in dynamics within forest climate sensitivity. Challenges remain in sourcing gridded, environmental covariates (e.g. Rapauch et al., 2009) at the moderately fine remote sensing scales (i.e. 20 m).

## Data Availability

Larger version of detailed images and figures can be accessed in a data repository at the following link: https://hie-pub.westernsydney.edu.au/projects/fmc_modelling/fmc_layers/other_layers/fmc_drought/

