## Supplementary for "Forests and woodlands resistant to drought revealed in remotely sensed foliar moisture content using probabilistic models"

### S1 FMC timeseries within landscapes

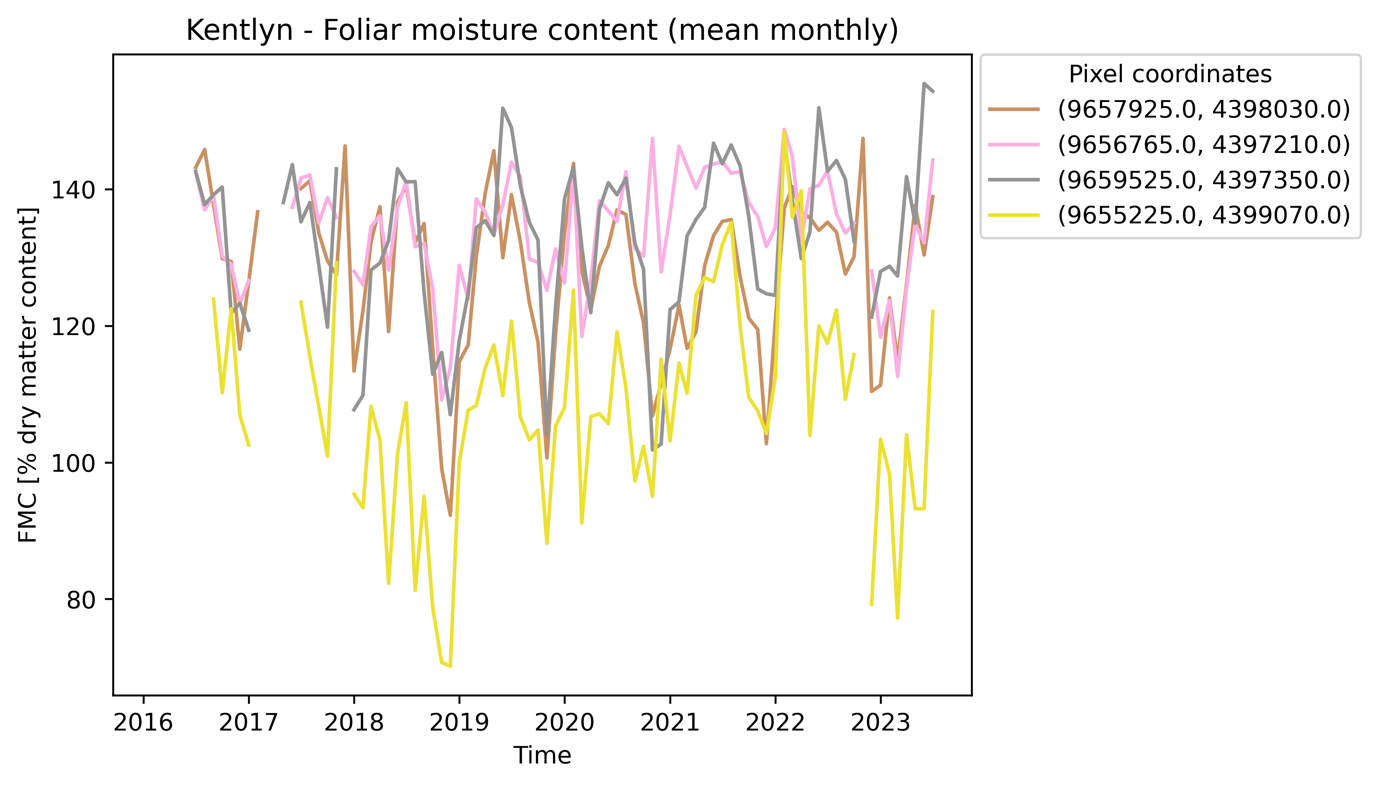

Figure S10. Mean monthly foliar moisture content timeseries of four random forest pixels in the landscape Kentlyn.

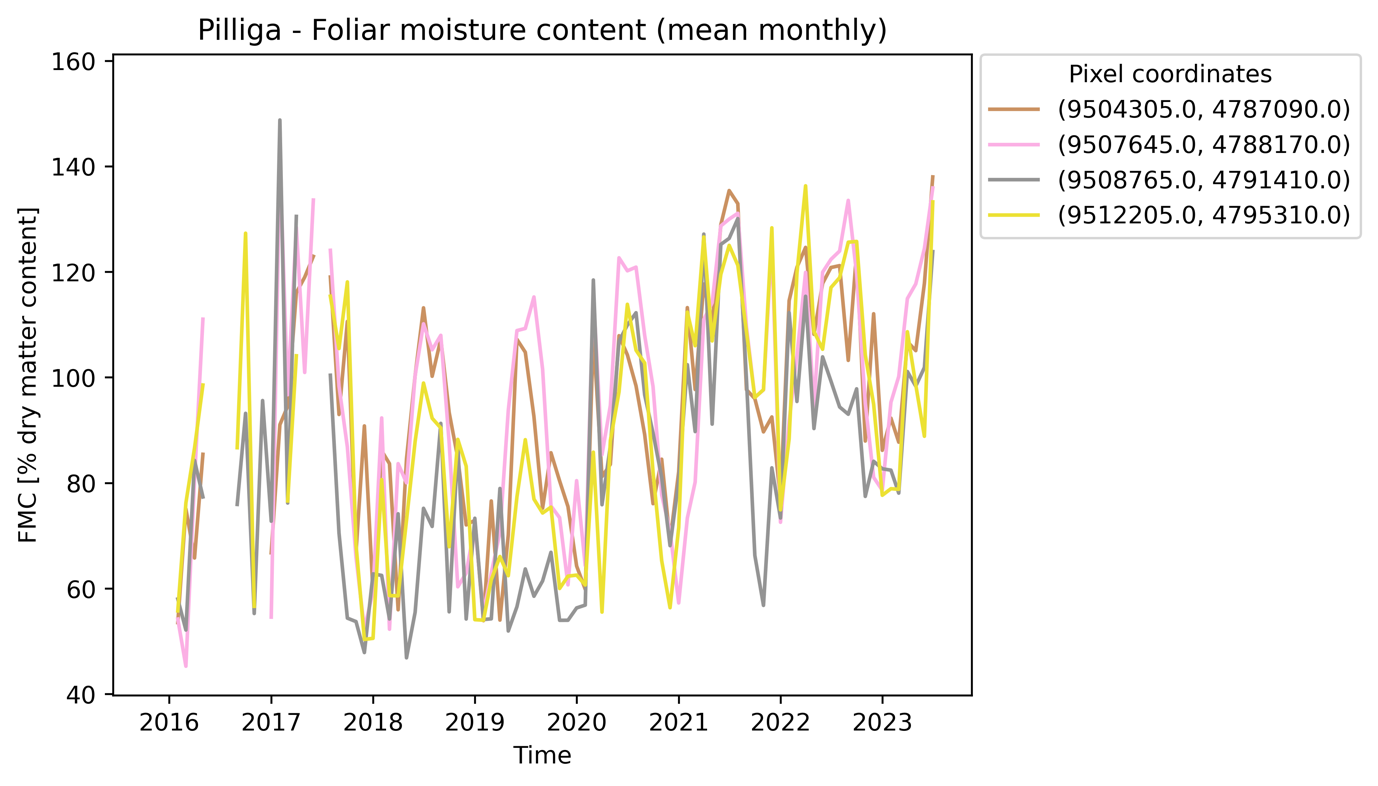

Figure S11. Mean monthly foliar moisture content timeseries of four random forest pixels in the landscape Pilliga.

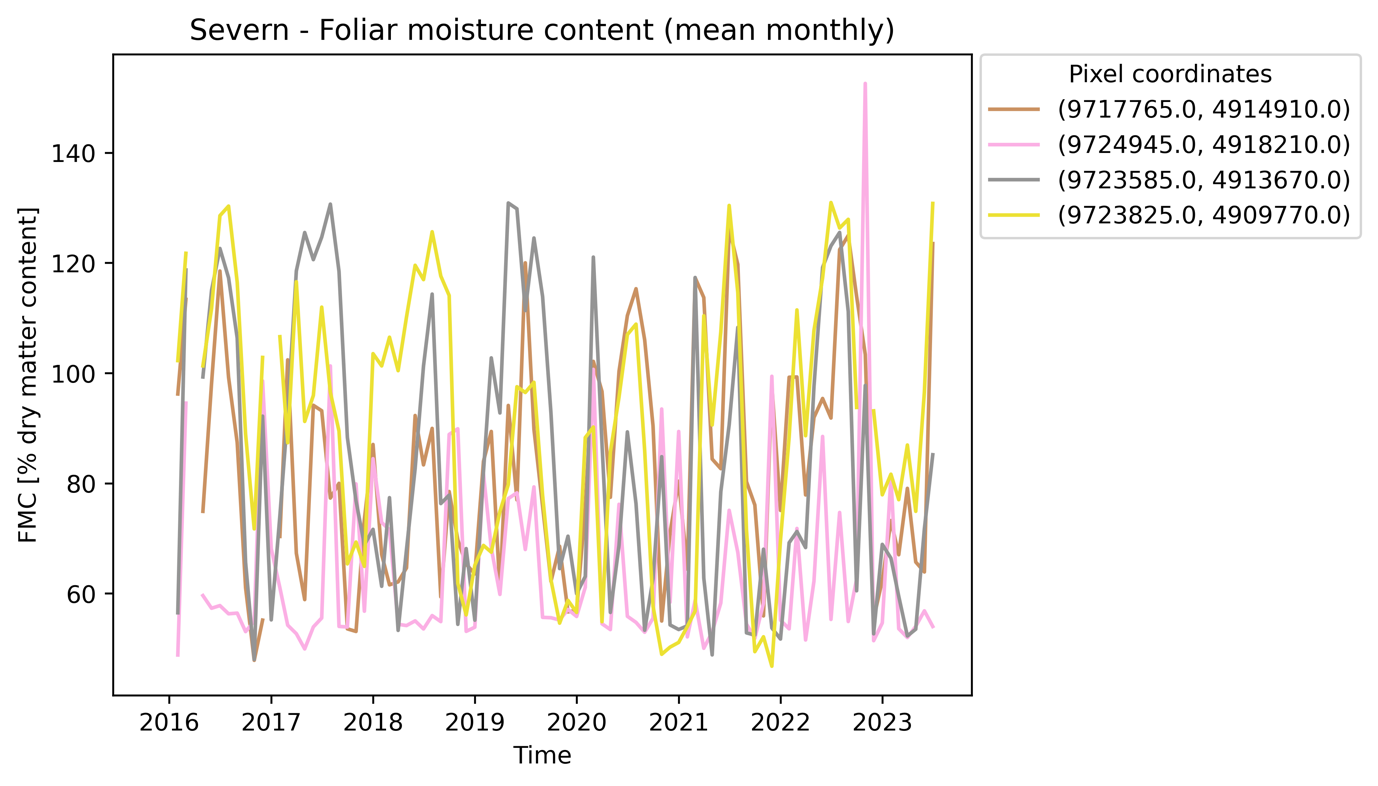

Figure S12. Mean monthly foliar moisture content timeseries of four random forest pixels in the landscape Severn.

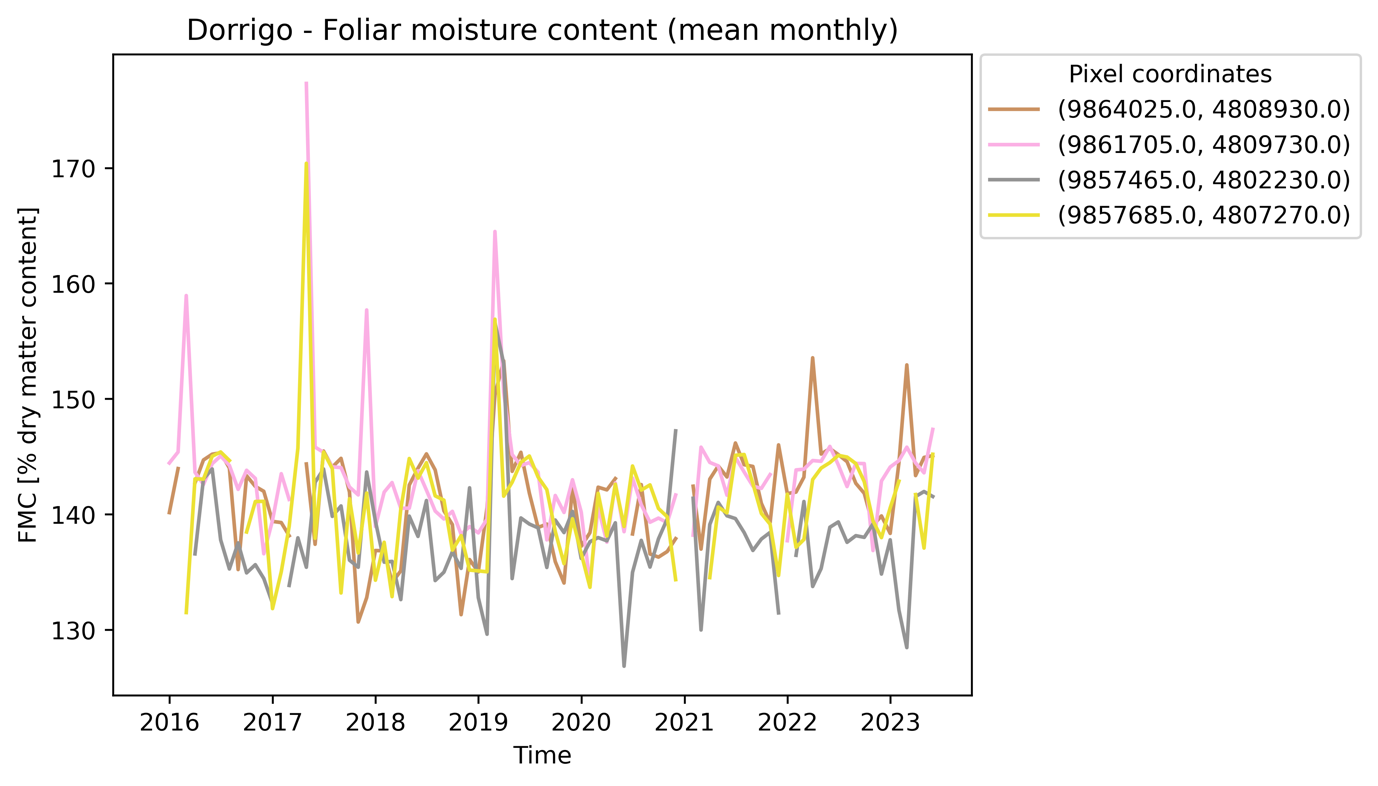

Figure S13. Mean monthly foliar moisture content timeseries of four random forest pixels in the landscape Dorrigo.

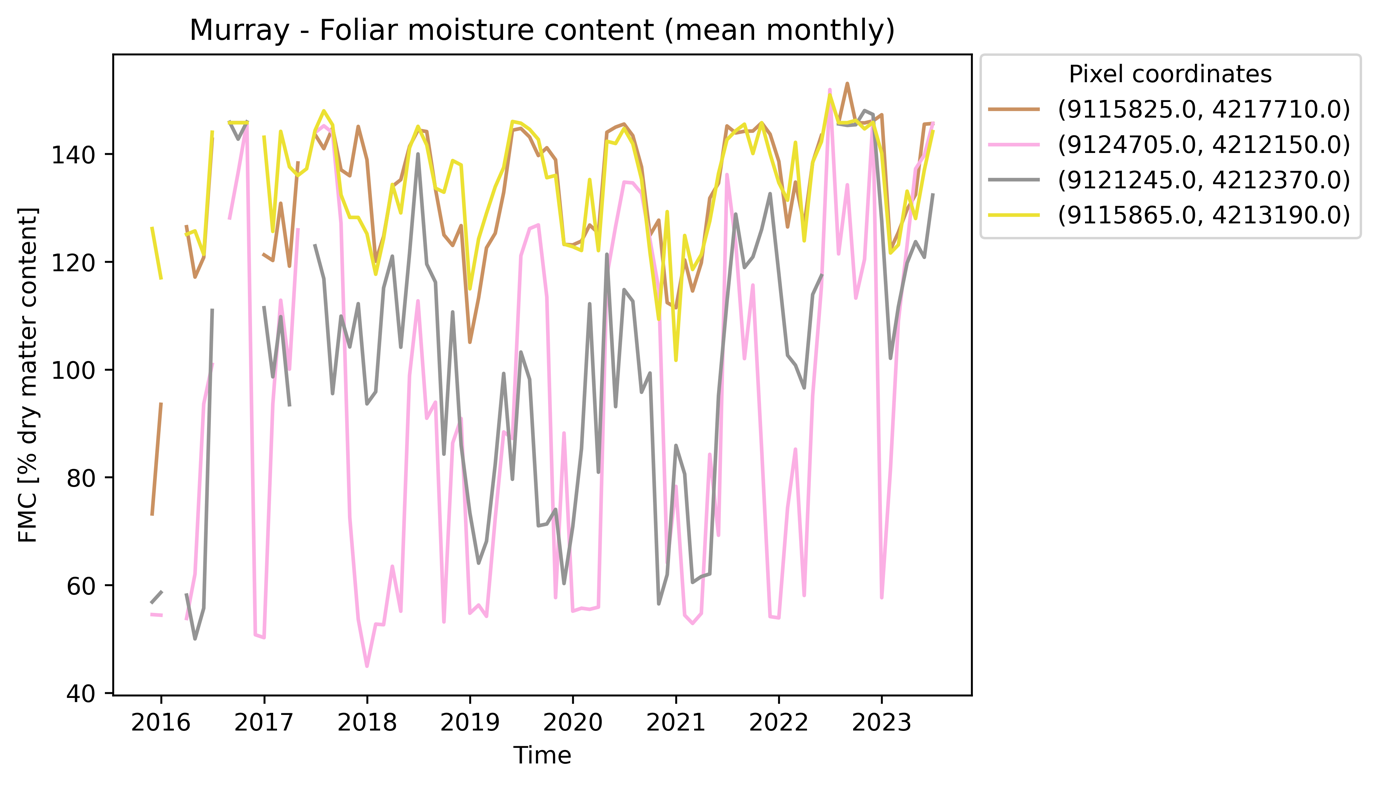

Figure S14. Mean monthly foliar moisture content timeseries of four random forest pixels in the landscape Murray.

### S2 FMC percentiles of landscapes

Table S1. Landscape means of absolute values of FMC percentiles used in the calculation of probabilities of FMC decline with drought.

|  |  | FMC | |
| --- | --- | --- | --- |
| Landscape | **Percentile** | **Mean** | **Std. dev.** |
| Kentlyn | 40 | 123.4 | 14.0 |
|  | 30 | 119.2 | 15.3 |
|  | 20 | 114. | 16.8 |
|  | 10 | 107.0 | 19.1 |
| Pilliga | 40 | 67.4 | 14.8 |
|  | 30 | 61.1 | 11.4 |
|  | 20 | 56.0 | 7.4 |
|  | 10 | 52.0 | 4.6 |
| Dorrigo | 40 | 141.1 | 5.3 |
|  | 30 | 139.7 | 6.1 |
|  | 20 | 138.0 | 6.9 |
|  | 10 | 135.5 | 7.8 |
| Severn | 40 | 66.9 | 16.8 |
|  | 30 | 61.9 | 13.9 |
|  | 20 | 57.4 | 10.9 |
|  | 10 | 53.0 | 7.7 |
| Murray | 40 | 116.8 | 22.8 |
|  | 30 | 109.5 | 25.6 |
|  | 20 | 102.0 | 27.3 |
|  | 10 | 93.6 | 28.1 |

### S3 FMC probability of decline in drought, and FPC

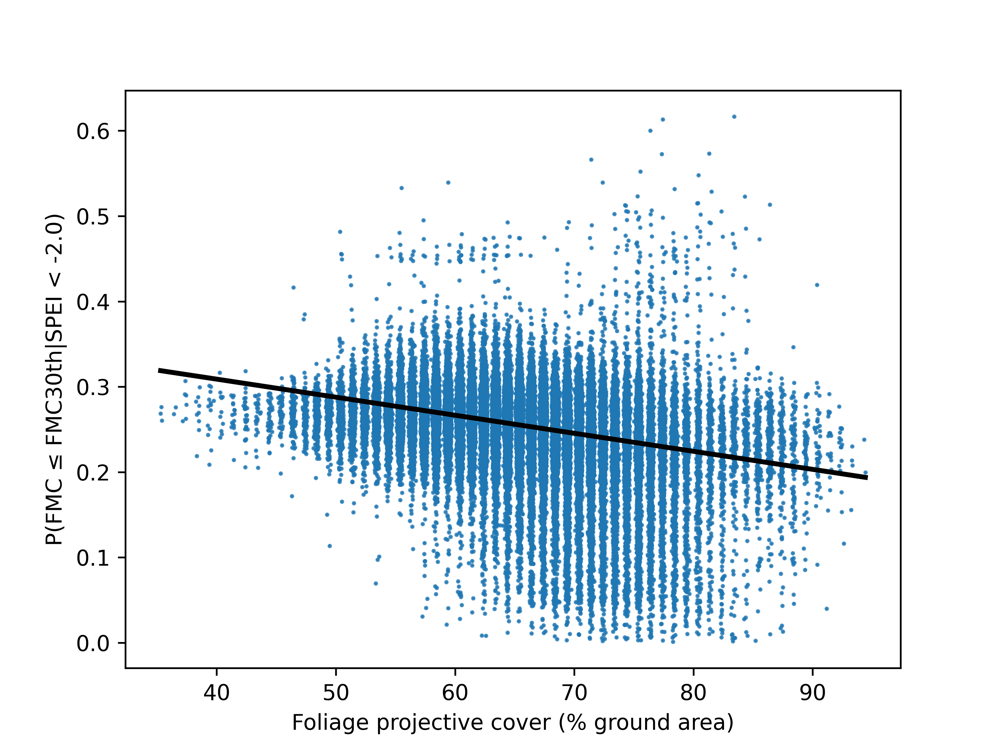

Figure S15. Probability of FMC decline to 30^th^ percentile in extreme drought against foliage projective cover at in the landscape at Kentlyn. Regression coefficients and results: slope=-0.002, intercept=0.39, R=-0.32, r^2^= 0.10, *P<*0.05. FPC is jittered by 0.4 for aesthetics because the product is estimated to nearest whole number.
